# A Physiologically Constrained Calibration Framework for Cardiovascular Models applied in Paediatric Sepsis

**DOI:** 10.64898/2026.02.10.704842

**Authors:** MT Cabeleira, S Ray, NC Ovenden, V Diaz-Zuccarini

## Abstract

Calibration of mechanistic cardiovascular models is a central barrier to their use in population analysis and patient-specific simulation, particularly in settings where key physiological variables are unobservable and multiple parameter combinations can reproduce the same haemodynamic targets. In this work, we present Embedded Gradient Descent (EGD), a calibration framework for ODE-based lumped-parameter cardiovascular models in which selected physiological parameters are promoted to dynamic states and driven toward prescribed targets through embedded controller equations. By exploiting the qualitative structure of the governing equations, EGD enforces physiologically consistent parameter–variable relationships, yielding unique calibrated solutions that are robust to initial conditions and scale efficiently with model complexity. The framework is demonstrated using a mechanistic cardiovascular model to generate virtual paediatric populations spanning normal physiology and two septic shock phenotypes (warm and cold shock), achieving low residual error across pressures, flows, and compartmental volumes. The resulting parameter distributions are consistent with known haemodynamic adaptations in paediatric sepsis, including alterations in vascular resistance, compliance, cardiac elastance, and effective blood volume. Importantly, persistent calibration residuals arise only when target combinations are structurally incompatible with the model, providing an explicit and interpretable diagnostic of feasibility limits rather than an optimisation failure. These results establish EGD as a general, scalable calibration strategy for mechanistic cardiovascular models and a practical foundation for virtual population generation and future patient-specific digital twin applications in critical care.

**NEW & NOTEWORTHY:** This study introduces a novel, embedded gradient descent calibration framework that enables scalable generation of mechanistically interpretable virtual populations of patients from ODE-based cardiovascular models. By treating parameter inference as a dynamical extension of the governing equations and calibrating directly against cycle-derived physiological targets, the method preserves physiologically meaningful parameter–variable relationships. Applied to paediatric sepsis, the framework reproduces warm and cold shock phenotypes while exposing infeasible target combinations, while providing efficient calibration and physiological insight.

## 1 INTRODUCTION

Lumped parameter models of the cardiovascular system (CVS) have been proven to be able to generate outputs that can mimic physiological behaviour with a high degree of precision [1, 2]. In this context, the CVS is usually conceptualized as an interconnected network of compartments, representing different parts of the CVS, mathematically formulated as systems of ordinary differential equations (ODEs) that exploit the fluid-electrical analogy, where pressures and flows of blood vessels are represented by their electrical analogues [3]. This modelling approach has found many applications in medical research over decades, being used in combination with more complex blood vessel models [4] or the in CircAdapt case where a complex heart model is informed by lumped parameter model of the CVS [5]. As stand-alone models, these can be used for the exploration of non-linear ventricular interactions [6, 7], orthostatic stress [8, 9] modelling or cardiopulmonary interactions [10, 11, 12, 13], where the crosstalk between the respiratory system and the CVS is explored. In Djoumessi et al. The addition of autonomic control to these models allows the study of the response to hypercapnia [14, 15, 16], obstructive sleep apnoea [17], cardiogenic shock [18] and other types of shock [19] and the response to fluid resuscitation technique [20].

Despite the proven versatility of these models in the medical field, efficient and fast parametrization is still a challenge, with the majority of the literature reusing parameters found in previous publications without providing much insight into the behavior of the model under different parameter regimes or under different physiological states [18, 21]. With the advent of digital patient twins and personalized medicine, the difficulty in parameterizing these models is hindering their translation to clinical practice in benefit of black box approaches. Some notable examples in literature that do provide solutions to the parametrization of ODE models usually also very much simplified [22, 23]. Some approaches use MCMC [24] and neural ODEs [25] to calibrate a model for acute circulatory failure or gradient descent, using the multishooting approach to calibrate a lung model [26]. When the models are of standard complexity, the literature provides insights into the parameter space and the most influential parameters via sensitivity analysis [17]. While these methods can be effective in specific settings, recent methodological guidelines for mechanistic cardiovascular modelling emphasise that calibration strategies should preserve physiological structure, maintain parameter interpretability, and explicitly expose identifiability and feasibility limits rather than relying on black-box optimisation approaches [27].

The problem in hand here is two-fold. Firstly, the most problematic aspect is the number of hidden variables (variables that are not measured) present in the standard-sized models, leading to the emergence of many redundant solutions when traditional optimization techniques are used. One possible approach here is to simplify the model and thus reduce the number of hidden variables or, if the number of observations is large enough, remove ambiguity completely. A good example is the work of Tannenbaum et al. [19] where the CVS was simplified into a 3 compartment model. This simplification is also tempting when traditional optimization techniques are used since these usually do not scale well with the number of parameters due to increasing computational requirements and dimensionality of the problem. Simplification is, however, not always possible as sometimes some hidden variables are fundamental for the the model to accurately reflect human physiology. A good example is the work of Albanese et al. [15] and Lu et al. [11] where capillary compartments are fundamental for modelling gas exchange between the lungs and the blood, despite capillary blood volume and pressure being impossible to measure.

Secondly, an aspect that is commonly overlooked in the literature is the effect total blood volume (TBV) has in the system, with most works focusing on the ‘traditional’ 75Kg adult male, or eliminating the need of volume variables all together by using ODE systems where pressure is the main primitive [19]. This is done because the TBV and its distribution across the system cannot be directly measured. The use of volume based ODE systems in scenarios where the TBV is not fixed generates another source of ambiguity to the optimization process and will also increase the number of parameters being estimated.

Septic shock provides a clinically relevant setting in which haemodynamic instability emerges from interacting changes in vascular tone, cardiac function, and TBV [28]. It is defined as a dysregulated host response to infection leading to life-threatening organ dysfunction [29], with septic shock representing a high-mortality subset characterised by severe circulatory, cellular, and metabolic abnormalities [30]. In paediatrics, recent international consensus criteria operationalise sepsis as suspected infection with a Phoenix Sepsis Score ≥ 2 and septic shock as sepsis with cardiovascular dysfunction (Phoenix cardiovascular subscore ≥ 1) [28], an approach that has been shown to improve the identification and prognostic stratification of paediatric sepsis compared with earlier IPSCC definitions [31]. In paediatric populations, sepsis remains a major cause of morbidity and mortality, accounting for more than 8% of paediatric intensive care unit admissions [32] and contributing substantially to global childhood mortality [33].

Clinically, septic shock is commonly classified into warm shock (WS) and cold shock (CS), reflecting distinct haemodynamic phenotypes [34], although clinical classification based on bedside signs alone has limited accuracy in identifying underlying haemodynamic states [32]. Warm shock is characterised by reduced systemic vascular resistance and preserved or elevated cardiac output, whereas cold shock is associated with increased vascular resistance and reduced cardiac output. Historically, early fluid resuscitation has been a central component of paediatric sepsis management [35]. However, subsequent evidence has demonstrated potential harm from aggressive fluid bolus therapy in specific settings [36], leading contemporary guidelines to recommend more cautious, context-dependent fluid strategies with early consideration of vasoactive support when shock persists [32].

Across both phenotypes, inflammatory-mediated increases in vascular permeability promote intravascular volume loss and redistribution into the interstitial space, reducing cardiac filling and arterial pressure. Compensatory autonomic responses attempt to restore perfusion through tachycardia, increased myocardial contractility, and vasoconstriction. In warm shock, however, these compensatory mechanisms may be ineffective due to uncontrolled peripheral vasodilation.

Figure 1 illustrates how distinct underlying mechanisms like volume redistribution, vascular tone dysregulation, or impaired myocardial contractility can give rise to overlapping haemodynamic signatures. Because TBV and its distribution cannot be measured directly, these similarities complicate diagnosis and treatment selection. This ambiguity motivates the use of mechanistic modelling approaches capable of explicitly representing blood volume, cardiac function, vascular resistance, and autonomic regulation within a unified physiological framework.

**Figure 1.**
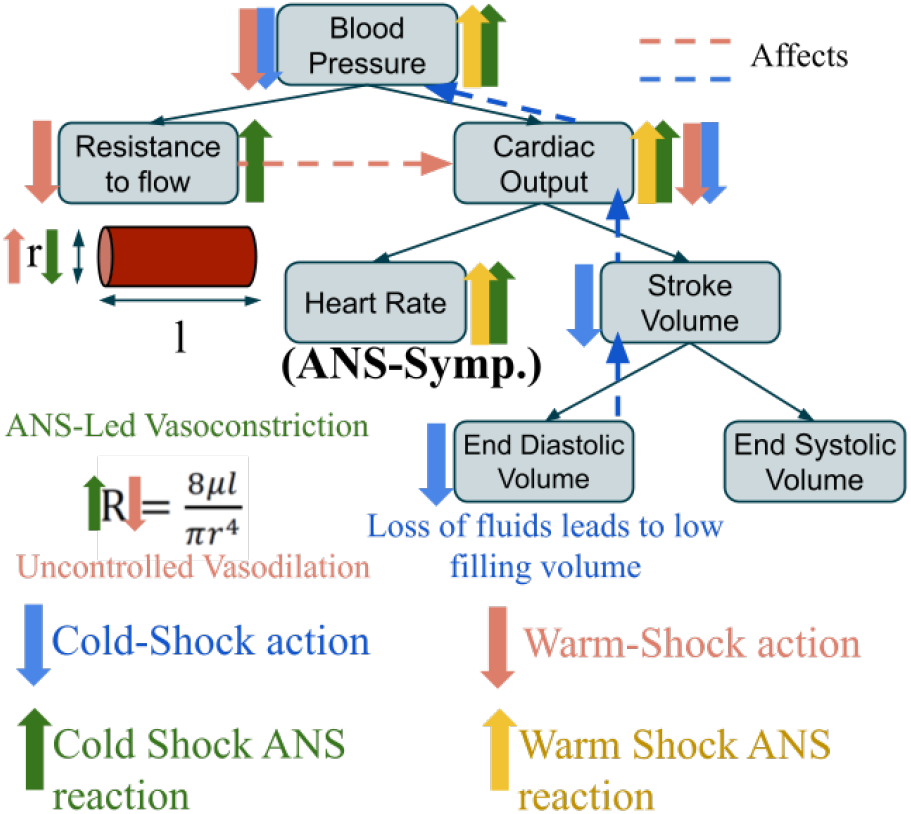
Schematic overview of the dominant haemodynamic mechanisms underlying warm and cold septic shock. Septic shock phenotypes are characterised by inflammatory-driven vascular permeability and blood volume redistribution, leading to reduced cardiac filling and hypotension, with compensatory autonomic responses acting on heart rate, contractility, and vascular resistance. In warm shock, peripheral vasodilation limits effective vasoconstrictive compensation, whereas cold shock is associated with elevated systemic vascular resistance.

The marked heterogeneity of sepsis, the coexistence of distinct haemodynamic shock phenotypes, and the need to account for variations in total blood volume make paediatric sepsis a particularly compelling setting for the development and testing of calibration techniques, as disease manifestations are more dynamic, age-dependent, and less well characterised than in adult populations, with current diagnostic and modelling approaches remaining limited [37]. In this work, we present a detailed explanation of our calibration approach entitled ‘embedded gradient descent’ EGD, which was initially introduced in our previous publication [10]. We further extend this algorithm to generate virtual populations that mirror the various clinical scenarios observed in paediatric sepsis patients. Specifically, we explain how first principles can be harnessed to identify the key model parameters and reveal their relationships with the observed variables and how to extend the number of observable variables for the optimization problem. We will also demonstrate its independence of initial conditions and ability to generate unambiguous model solutions.

## 2 MATERIALS AND METHODS

Here, we first describe the clinical data routinely available in a paediatric intensive care setting and identify the subset of observables relevant to the proposed model. We then introduce the cardiovascular model formulation and its governing equations. Finally, we present the embedded gradient descent calibration strategy and describe how physiological constraints are leveraged to generate patient-specific targets and virtual populations. All symbols, auxiliary operators, and time variables used in paper are summarised in the nomenclature in Appendix A.1.

### 2.1 Data available

A paediatric patient in a standard PICU is usually monitored by a combination of medical devices, depending on the condition and its severity. While continuous waveform recordings are technically feasible and available in some centres, their routine clinical use is limited by data volume, storage, and downstream usability constraints. As a result, many clinical data integration systems prioritise the recording of lower-frequency, trend-level physiological data, with sampling intervals on the order of seconds. The variables routinely recorded from these patients that are relevant for our model are:

- Systemic Arterial Blood Pressure (ABP) - 3 available pressures: systolic, diastolic and mean systemic blood pressures.
- Heart Rate (HR)
- Central Venous Pressure (CVP) - value of the mean blood pressure in the systemic venous system.
- Cardiac Output (CO) - A measure of the total amount of blood that flows through the heart over a minute. Commonly measured indirectly, including via echocardiography or less commonly, via tracer dilutional techniques based on the Fick principle.
- Pulmonary arterial Pressure (PAP) - usually extrapolated via echo-cardiography or directly measured using a Swan-Ganz catheter, although the use of this is very rare. In addition, the pulmonary capillary wedge pressure (PCWP), can also be obtained during balloon occlusion.

Patient demographics, including; age, height, weight, sex, are also collected and used to contextualize the values of the other variables collected or to estimate other parameters like TBV.

#### 2.1.1 Physiological priors derived from clinical practice

Clinical monitoring in the intensive care setting provides access to only a limited subset of cardiovascular variables, whereas many physiologically relevant quantities, including blood volume distribution, regional pressure levels, and microcirculatory pressures, are not directly observable in routine practice. To address this gap, we introduce a set of physiological priors derived from established clinical knowledge and population-level studies. These priors represent plausible reference values and relationships that constrain otherwise unobserved quantities. Their role is to provide physiologically consistent bounds and relationships that will later be used to contextualize the available data and to support the construction of individualized cardiovascular representations.

#### Capillary Pressures

While capillary pressure is not directly measurable in routine clinical settings, physiologically plausible target values can be inferred by considering the expected pressure gradients along the vascular tree as blood flows from the arterial to the venous circulation. This allows a pressure drop between the arterial and capillary beds to be prescribed, from which a corresponding range of capillary pressures can be derived.

#### Total Blood Volume (TBV)

The cardiovascular system is commonly treated as conserving TBV over short time scales under stable conditions. However, the absolute blood volume of an individual is not directly measurable, nor is its distribution across the circulation. In clinical practice, the partial solution found for this is the estimation of TBV involving the use of different formulae for different patient populations using the patients sex, height and weight. For example using the Nadler’s formulae where height, weight are used to estimate TBV. Other methods use body weight alone as in Ranaa et al. [38] or in Raes et al. [39] where a known amount of albumin marked with iodine is injected and allowed to homogenize with the blood, the TBV is then estimated by calculating the diffused albumin concentration.

To estimate a possible value for the average blood volume within different regions of the circulation, the values of Table 1 were used, together with the estimation of the TBV. These assumptions provided a plausible value for every volume variable in the equation system, with the downside of considering every patient to be ‘normal’ in terms of the distribution of volume.

**Table 1.** Average blood volumes on different parts of the circulatory system, assuming a TBV of 700mL. Extracted from [40].

| Cardiovascular Region | Volumes |  |
| --- | --- | --- |
|  | mL | % |
| Left Heart | 24.5 | 3.5 |
| Right Heart | 24.5 | 3.5 |
| Systemic arteries (Aorta) | 91 | 13 |
| Systemic arterioles & capillaries | 49 | 7 |
| Systemic Veins | 448 | 64 |
| Pulmonary Arteries | 26.6 | 3.8 |
| Pulmonary Capillaries | 9.8 | 1.4 |
| Pulmonary Veins | 26.6 | 3.8 |

#### Flow Constraint

Over sufficiently long time scales, the average blood flow through the cardiovascular system is governed by the CO, which represents the volume of blood pumped by the heart per unit time. As a result, the mean flow supplying and draining the major regions of the circulation can be assumed to be equal to CO.

#### 2.1.2. Physiological target ranges for paediatric sepsis

To characterise physiologically plausible ranges for the sepsis scenarios considered in this study, the clinical literature was surveyed and the resulting target values are summarised in Table 2.

**Table 2.**
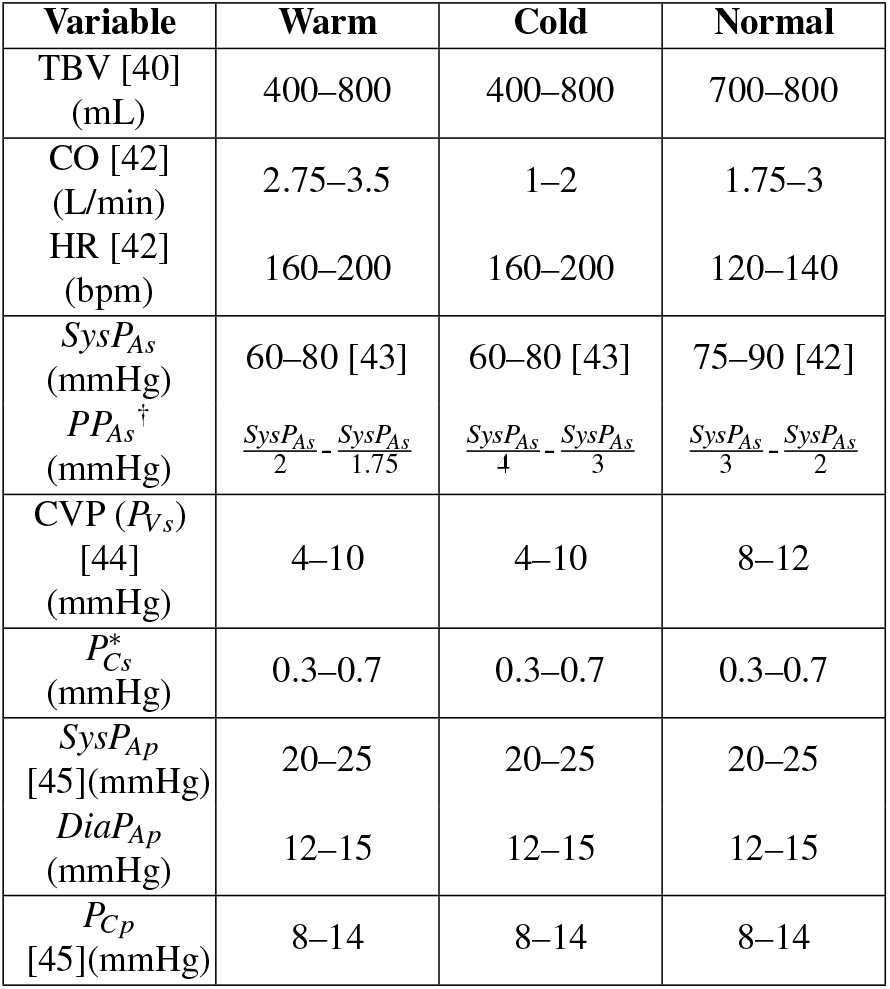
Target ranges used for calibration under warm shock, cold shock, and normal physiology. *As the capillary pressure of the systemic system is impossible to measure, we have imposed a % drop in pressure from the diastolic value of the Systemic arterial pressure and the value of CVP. † The targets for the Pulse Pressure of the systemic arterial blood pressure is calculated according to the equation extracted from Ranjit et al. [43] where these ratios were found to be descriptive of sepsis cases.

| Variable | Warm | Cold | Normal |
| --- | --- | --- | --- |
| TBV [40] (mL) | 400–800 | 400–800 | 700–800 |
| CO [42] (L/min) | 2.75–3.5 | 1–2 | 1.75–3 |
| HR [42] (bpm) | 160–200 | 160–200 | 120–140 |
| $SysP_{As}$ (mmHg) | 60–80 [43] | 60–80 [43] | 75–90 [42] |
| $PP_{As}^{\dagger}$ (mmHg) | $\frac{SysP_{As}}{2} - \frac{SysP_{As}}{1.75}$ | $\frac{SysP_{As}}{4} - \frac{SysP_{As}}{3}$ | $\frac{SysP_{As}}{3} - \frac{SysP_{As}}{2}$ |
| CVP ($P_{Vs}$) [44] (mmHg) | 4–10 | 4–10 | 8–12 |
| $P_{Cs}^*$ (mmHg) | 0.3–0.7 | 0.3–0.7 | 0.3–0.7 |
| $SysP_{Ap}$ [45] (mmHg) | 20–25 | 20–25 | 20–25 |
| $DiaP_{Ap}$ (mmHg) | 12–15 | 12–15 | 12–15 |
| $P_{Cp}$ [45] (mmHg) | 8–14 | 8–14 | 8–14 |

#### TBV

The reference subject was a 12 month old child with a body surface area of 0.5 *m*^2^ and therefore would have, in a healthy scenario 700-800 mL of TBV. As shock can happen in normovolemia and hypovolemia, the TBV of the shock cases was set between 400-800 *mL*.

#### Cardiac output & HR

The cardiac index that is considered normal by Brierley et al. [41] is between 3.5 and 6.0, thus if WS is characterized by increased CO, it would have CI between 5.5 and 7 and CS having low CO would have a CI of 2-4. These limits were chosen so that there would be some overlap with the ‘normal’ class whilst allowing for the exploration of more extreme cases as well. According to sepsis guidelines, a heart rate above 160 is considered clinically concerning, while values exceeding 190 are regarded as an indicator of sepsis (in the context of suspected or proven infection) [42].

#### Systemic blood pressures

The systolic blood pressure *SysP*_*As*_ is considered clinically concerning when below 70 [42] to 75 mmHg [46] and the normal values were found to be between 75-90 mmHg [42]. As both shocks can have normal to low systolic blood pressure [43], the limits for these were set to 60-80 mmHg. Systemic pulse pressure (*PP*_*As*_), defined as the difference between systolic and diastolic pressures in the systemic arteries, is considered clinically concerning when *PP*_*As*_ *> SysP*_*As*_*/*2. In this context, warm shock is typically associated with normal to elevated *PP*_*As*_, whereas cold shock is characterised by a narrowed *PP*_*As*_ [43]. For this we set the limits for PP as fractions of *SysP*_*As*_ with the WS always being *> SysP*_*As*_*/*2 and the CS *< SysP*_*As*_*/*3. These pulse-pressure targets are inherently heuristic and may break down at elevated heart rates, where pressure-based indices no longer reliably reflect cardiac performance, as discussed by Tibby et al. [47].

After fluid resuscitation therapy, the values for CVP that lead to the best outcomes are observed to be between 8-12mmHg [44] and thus these were the limits set for ‘normal’. The shock cases having normal to low CVP, were set to 4-10 mmHg. As the pressure at the systemic capillaries is difficult to measure, the targets for these were set as a % pressure drop between the diastolic pressure and the CVP. The range for this % pressure drop was set to 0.3 to 0.7 in all cases to simulate both high and low capillary pressures.

#### Pulmonary pressures

On the pulmonary system, sepsis specific studies where the pulmonary parameters are mentioned were not found, with the literature focusing more on pulmonary hypertension, with the guidelines stating that systolic Pulmonary Arterial Blood Pressure sPABP *>* 35mmHg [48, 49] and mean pulmonary pressure *>* 20 mmHg [45] and Pulmonary Capillary Wedge Pressure PCWP target between 12-14 mmHg [45]. With this information we set the sPABP to be ‘normal’ in all subjects (20-25 mmHg) and the diastolic pressure to be 12-15 mmHg to allow some pressure drop before the capillaries. To these we set a target capillary pressure to be 1-4 mmHg lower than the diastolic pressure, generating capillary pressures with a range between 8-14 mmHg.

### 2.2 Cardiovascular system model

The physiological target ranges and population constraints defined above specify the haemodynamic conditions that the model must reproduce. We now introduce the cardiovascular system model and its governing equations, which provide the mechanistic basis for subsequent calibration.

The cardiovascular system model used in this work is illustrated in Figure 2. It represents the heart and circulation as a lumped-parameter network composed of interconnected compartments corresponding to the major functional regions of the cardiovascular system. Each compartment represents a spatially aggregated vascular or cardiac region and is characterised by a pressure, a volume, and associated inflow and outflow rates. The compartments are connected through resistive elements that govern blood transport between regions, while capacitive elements account for vascular and cardiac compliance.

**Figure 2.**
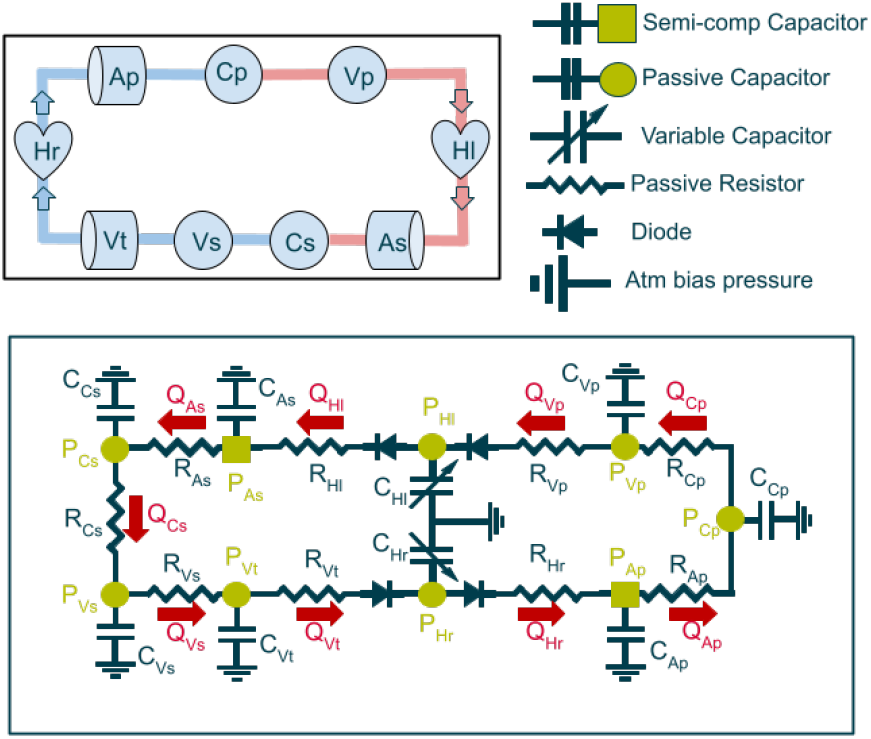
Schematic representation of the cardiovascular model. **Top:** High-level block diagram showing the major compartments of the systemic and pulmonary circulations, including the left and right heart chambers (Hl, Hr), arteries (As, Ap), capillaries (Cs, Cp), veins (Vs, Vp), and thoracic veins (Vt). **Bottom:** Full circuit representation of the model, where each compartment is represented by a compliance element and connected to neighbouring compartments through resistances that govern inflow and outflow.

This formulation results in a closed-loop circulation comprising nine compartments, and therefore nine state pressures and volumes, together with the flows that couple them. The model structure follows the same modelling strategy presented in [10], where the approach is described in greater detail. In the present work only the heart and circulation are represented, as these are the primary focus of the analysis.

The governing equations that define the pressure, volume, and flow dynamics of each compartment are introduced in Section 2.2.1. The heart chambers require additional treatment to capture their cyclic contraction and relaxation, which is addressed through time-varying elastance models and a dedicated heart-rate formulation described in Section 2.2.2. Finally, a set of auxiliary equations is introduced to compute cycle-based quantities such as extrema, averages, and integrals of physiological variables, which are required for analysis and calibration and are described in Section 2.2.3.

#### 2.2.1 Model equations

The model shown in Figure 2 consists of nine compartments, each of which is described using the same generic compartment formulation illustrated in Figure 3. The equation system that describes the generic compartment is:

**Figure 3.**
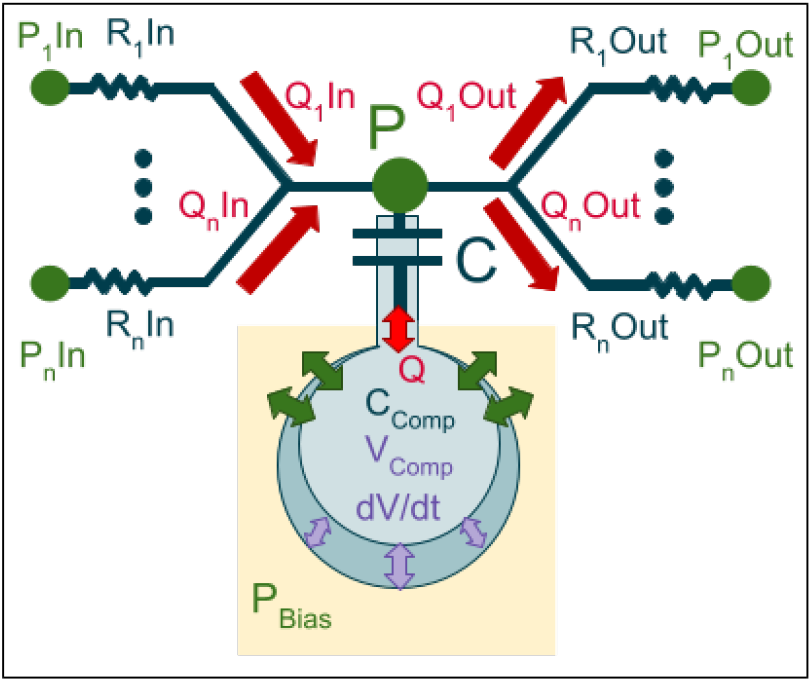
Generic representation of a compartment in the cardiovascular model. Each compartment is connected to its neighbours through a set of resistors, characterised by inlet and outlet resistances *R*_*in*_ and *R*_*out*_, which govern the inflow *Q*_*in*_ and outflow *Q*_*out*_ according to the pressure differences with neighbouring compartments (*P*_*in*_,*P*_*out*_). The central capacitor represents the compartment’s compliance *C*, which stores a volume of fluid and determines the local pressure *P*. Together, these elements define the fundamental pressure/volume/flow relationships used throughout the full model.

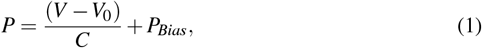

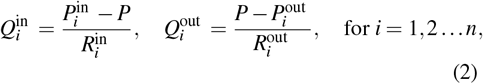

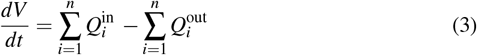

In Equation 1, *P* denotes the pressure within the compartment, *V* the compartmental blood volume, *V*_0_ the unstressed volume, *C* the compliance, and *P*_Bias_ an additive pressure offset accounting for external reference pressures. Equation 2 defines the inflow and outflow rates 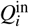 and 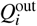 between the compartment and its *n* neighbouring compartments, driven by the pressure differences 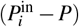 and 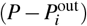 across the corresponding inlet and outlet resistances 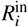 and 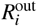. The temporal evolution of the compartmental volume is then given by Equation 3, which enforces mass conservation by equating the rate of change of volume to the difference between total inflow and total outflow.

The resistors at the inlet and outlet of the heart compartments are modelled as diodes to prevent backflow of blood, ensuring unidirectional flow during the cardiac cycle.

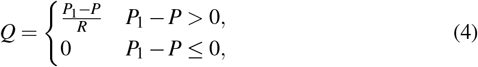

#### 2.2.2 Cycles: Heart rate modelling

The heart functions as a muscle that operates in a periodic cycle of contraction (systole) and relaxation (diastole). Consequently, the compliances representing the heart chambers also have to be specifically tailored to account for the cyclic nature of cardiac function. In literature, the most common approach is to modulate the value of compliance using elastance models (ℰ = 1*/C*). We use the variable-elastance model adapted from Heldt et al. [8] written as:

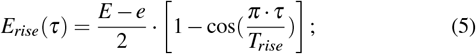

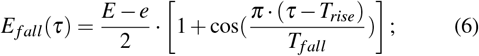

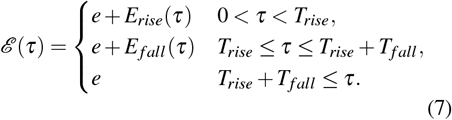

where *E* and *e* represent the maximum (systolic) and minimum (diastolic) elastances of the chamber, respectively, *T*_*rise*_ and *T*_*fall*_ define the durations of the systolic contraction and relaxation phases, ℰ (*τ*) represents the resulting time-varying elastance over the cardiac cycle. *HR* is the heart rate, *HC* the duration of a single heart cycle.

The heart elastance equations (7) require a time variable *τ* that represents the elapsed time within the current cardiac cycle rather than absolute simulation time. The duration of each cycle is determined from the heart rate according to

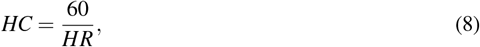

where *HR* denotes the heart rate in beats per minute and *HC* the resulting duration of a single cardiac cycle.

To ensure that changes in heart rate affect only subsequent cardiac cycles and do not perturb the ongoing one, the model employs an event-triggered cycle mechanism. As illustrated in Fig. 4, the trigger equation (9) schedules the onset time of the next cardiac cycle, stored in the variable *T*_trig_, based on the current cycle duration. When the trigger condition is met, the timer equation (10) resets the intra-cycle time *τ* to zero, after which it advances linearly until the next trigger event. Reset operations are implemented as fast relaxations over a single numerical integration step of duration Δ*t*, which corresponds to the fixed time step of the solver. This choice avoids explicit hybrid events while ensuring that cycle resets occur effectively instantaneously at cardiac boundaries. Throughout, *t* denotes absolute simulation time, while *τ* represents the elapsed time within the current cardiac cycle and provides the phase variable used in the elastance model.

**Figure 4.**
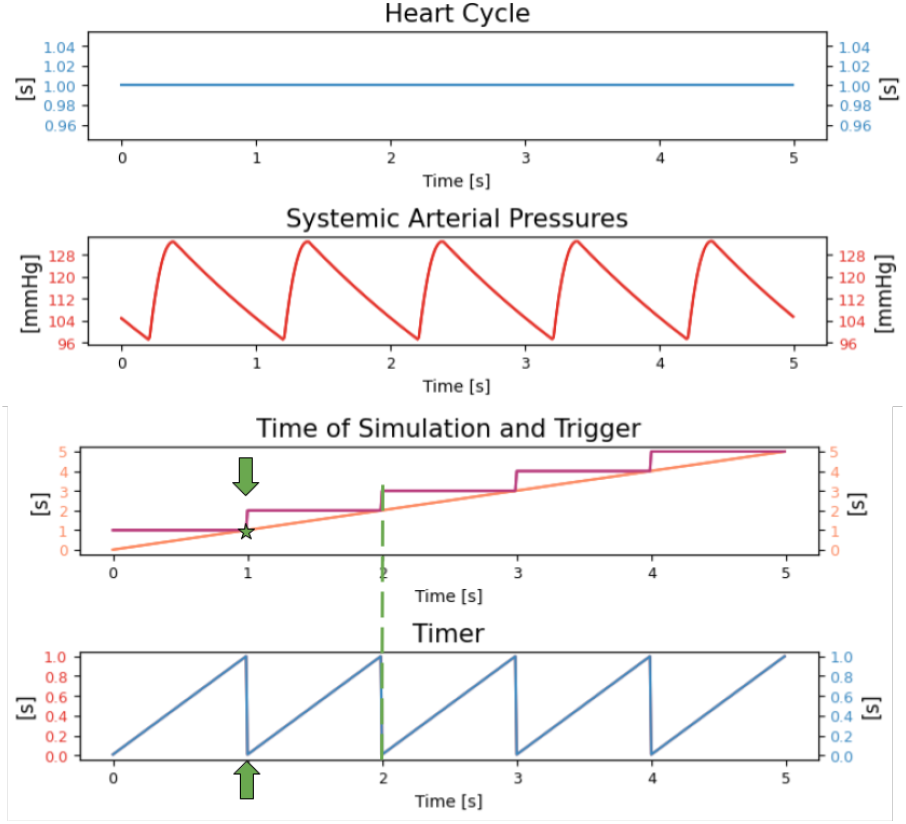
Schematic representation of the mechanism used to enable variable heart rates within the simulation framework. Top panel: prescribed heart-cycle duration. Second panel: resulting systemic arterial pressure waveform over consecutive cardiac cycles. Third panel: absolute simulation time (orange) and discrete trigger time (purple); when the trigger value coincides with the absolute time, it is advanced to the next scheduled trigger instant. Bottom panel: cycle timer, which resets to zero upon each trigger update and evolves linearly until the next cardiac event.

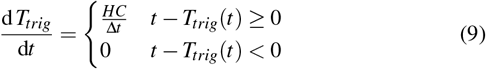

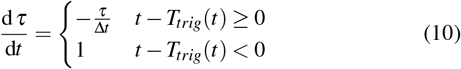

#### 2.2.3 Calculation of max, min, average and integration over cycle

The trigger mechanism described earlier is also used to compute cycle-based physiological quantities required during simulation and calibration (Section 2.3). The trigger resets the evaluation window for maxima, minima, integrals, and other beat-to-beat metrics. The maximum and minimum of a variable *Y* are obtained using Eqs. 11, 12, which reset their values at each trigger and track the extremes until the next cycle.

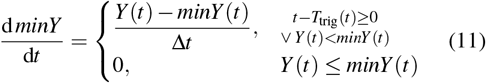

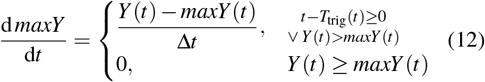

Stroke Volume is computed via the integral operator in Eq. 13, which accumulates flow until the next trigger event and resets thereafter.

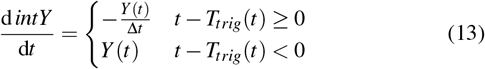

Because the quantity of interest corresponds to the value accumulated over the previous cardiac cycle, a supporting keeper equation (Eq. 14) is used to store the pre-reset value at each trigger event and hold it constant throughout the subsequent cycle

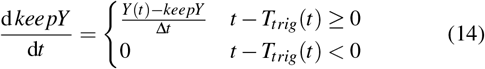

Finally, moving averages are computed continuously using Eq. 15, where the averaging period (*T*_avg_) defines the temporal window of interest. Together, these equations enable online computation of cycle-based quantities required for real-time calibration and analysis.

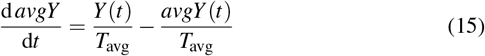

Here, *Y* (*t*) denotes an arbitrary scalar physiological variable of interest (e.g., pressure, flow, or volume), and *MinY, MaxY, IntY, AvgY*, and *KeepY* denote auxiliary state variables used to compute cycle-based statistics.

These operators allow cycle-resolved quantities to be computed online during integration, enabling direct comparison between simulated outputs and clinically defined target metrics used during model calibration.

#### 2.2.4 Final system of equations

The complete system of ordinary differential equations is obtained by instantiating the generic model equations defined above for each cardiovascular compartment using the unified specification provided in Table 3. This table encodes, the compliance and elastance elements associated with each compartment, the resistive connections governing inter-compartmental flows, and the corresponding volume balance relationships, fully defining the network topology illustrated in Fig. 2.

**Table 3.**
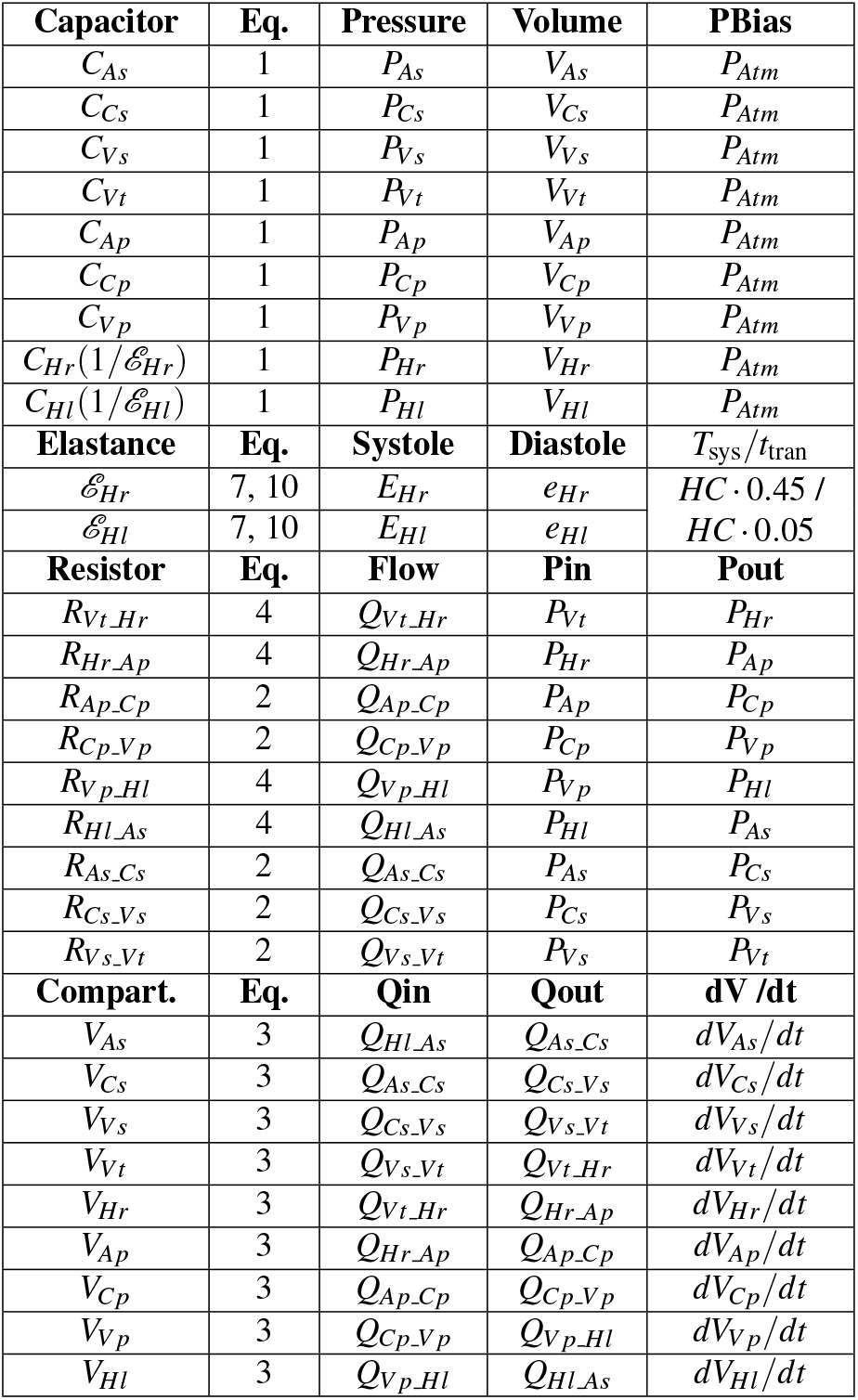
Unified specification of the cardiovascular model. The table defines all compartmental compliance and elastance elements, inter-compartmental resistive connections, and volume balance relationships required to instantiate the full closed-loop system shown in Fig. 2.

For each compartment, the pressure/volume relationship is constructed from Eq. 1, with heart chambers additionally governed by the time-varying elastance formulation in Eq. 7. Inter-compartmental flows are generated from the specified resistive links using either the linear resistance formulation (Eq. 2) or the diode model (Eq. 4), and compartmental volumes evolve according to mass conservation as expressed in Eq. 3. Together, these instantiated equations yield a closed-loop cardiovascular system with coupled pressure, volume, and flow dynamics.

This formulation results in a system with 27 primary state variables, comprising 9 compartmental pressures, 9 compartmental volumes, and 9 inter-compartmental flow variables. The model includes 29 physiological parameters, consisting of resistances, compliances (or elastances), and unstressed volumes (*R, C*/*E, V*_0_). Two additional parameters arise from the heart compartments, which are characterised by maximum and minimum elastances rather than fixed compliances.

### 2.3 Model Calibration

Calibrating the cardiovascular model to the population requires identifying parameter values that reproduce the characteristic pressures, volumes and flows of that population while respecting the intrinsic relationships encoded in the governing equations. Because each variable is influenced simultaneously by all compliances, resistances and volumes, changes in any parameter propagate throughout the entire system. This coupling makes direct parameter assignment difficult and undermines traditional optimization methods, which adjust parameters independently and ignore the monotonic interactions built into the model. By operating outside the model’s dynamical structure, these approaches attempt to match target pressures or flows without using the natural correlations encoded in the equations. Because several parameters can influence the same variable in similar ways, they cannot reliably determine which adjustments are appropriate, often require extensive computational search to find acceptable solutions, and frequently return multiple distinct parameter sets that satisfy the same targets. As the number of variables increases, these methods scale poorly, becoming increasingly resource intensive.

To address these issues, we introduce an embedded calibration strategy entitled Embedded Gradient Descent (EGD) described in Section 2.3.1. Here, rather than keeping the calibration parameters as fixed constants, we add dedicated differential equations that govern their evolution, each constructed according to the qualitative correlations between parameters and variables. These new equations operate alongside the physiological ODEs and drive the parameters in the direction required to reduce their respective errors, continuously ‘nudging’ the system toward the prescribed targets.

Because the EGD framework introduces parameter dynamics directly into the model, the corresponding correction equations must be tailored to the physiology and cannot be assigned arbitrarily. Section 2.3.2 explains how these equations were selected by leveraging the structure of the governing ODEs, the qualitative correlations between parameters and variables, and established physiological principles. This stands in contrast to traditional plug-and-play optimization methods, which adjust parameters without regard for how those adjustments align with the underlying system behavior.

An overview of the Embedded Gradient Descent (EGD) calibration framework is shown in Fig. 5. The diagram summarises how clinical observables are mapped onto model variables within the cardiovascular network, how the governing physiological equations and embedded controllers interact and how parameter adaptation is driven by target errors to steer the system toward a physiologically consistent steady state. The formal definition of the calibration dynamics is introduced in the following subsection.

**Figure 5.**
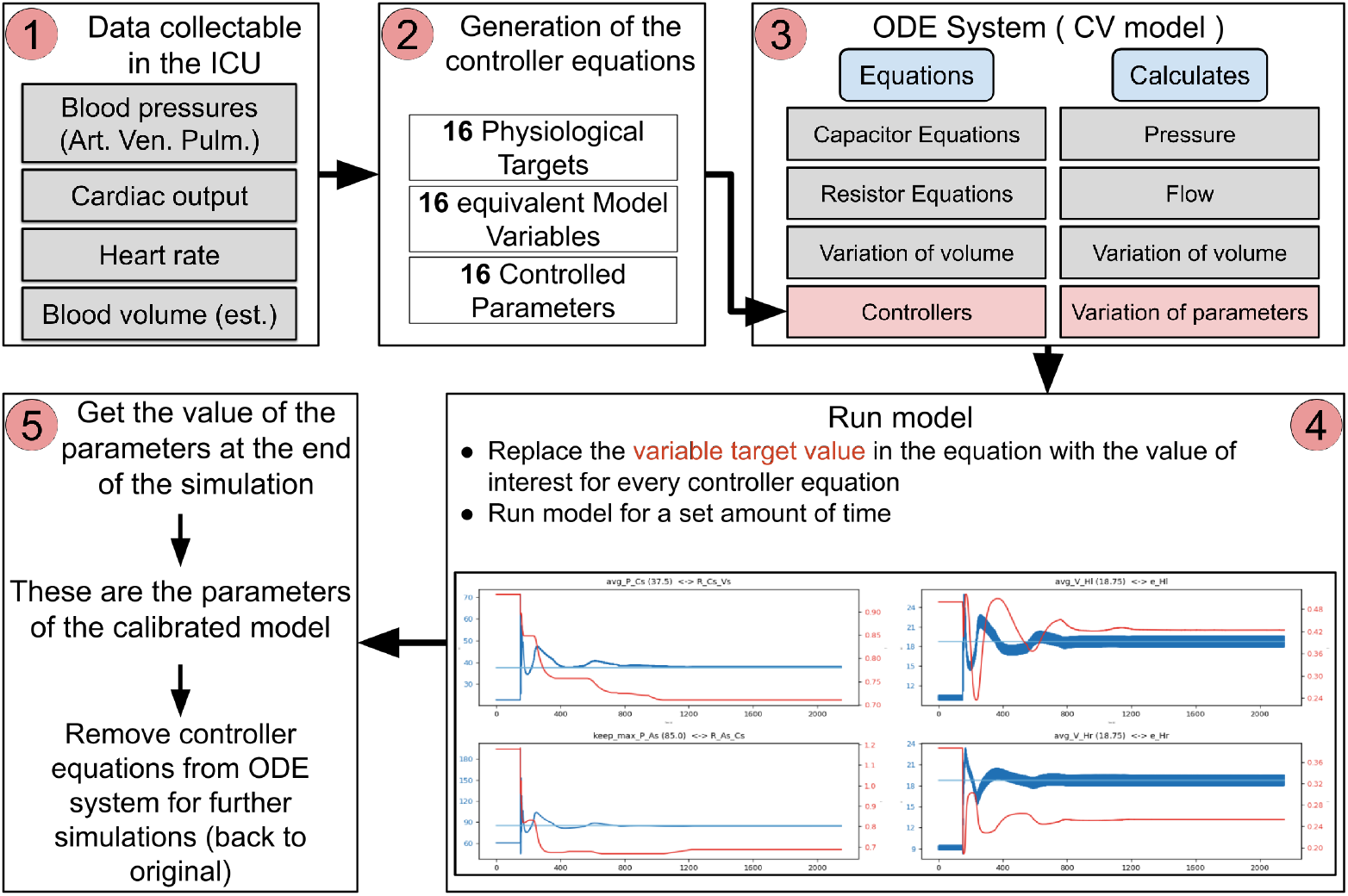
Workflow of the calibration methodology using Embedded Gradient Descent (EGD). (1) Physiological quantities that can be measured in the ICU are the calibration targets. (2) These clinical targets are mapped onto the corresponding model variables and the parameters that influence them, establishing the link between observable data and the internal states of the cardiovascular model. (3) The full model is then assembled by combining the governing physiological equations (capacitor, resistor, and volume-balance relations) with the EGD controller equations that prescribe how selected parameters evolve in response to their relative errors. (4) The model is run with all controllers active, allowing pressures, flows, volumes, and parameters to co-evolve until the system converges toward the prescribed target state. (5) Once equilibrium is reached, the resulting parameter set is extracted, providing a calibrated, physiologically consistent representation of the patient or target population.

#### 2.3.1 Embedded Gradient Descent

Careful observation of eq. 1-3 allows us to extract the correlation between all the parameters and variables, as shown in Table 4. These correlations describe monotonic qualitative effects and do not imply linearity or exclusivity, but they are sufficient to define the sign of each calibration controller.

**Table 4.**
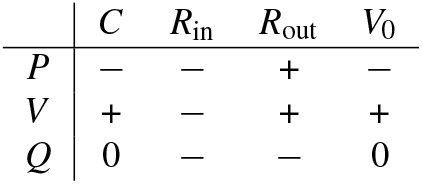
Qualitative Correlation Matrix for a Generic Compartment where the model variables of Pressure, Volume and Flow are compared to the parameters for Compliance, Resistance and unstressed volume (*V*_0_)

This table tells us that the task of increasing pressure in a compartment can be done either via increasing the outlet resistances (*R*_*out*_) or decreasing the inlet resistances (*R*_*in*_), compliance (*C*) or unstressed volumes (*V*_0_). Volume can be increased by increasing (*C*), *R*_*out*_ and *V*_0_ or decreasing *R*_*in*_. Reducing all resistances will increase the flow through the system and decreasing the compliance will also indirectly increase flow through as less fluid is retained in the compartment.

Knowledge of these relations between the parameters and the variables can be leveraged to create an extra set of differential equations that will drive the model to settle into physiologically relevant zones where the values for pressure, volume and flow generated by the model can be actively targeted. This can be done by reinterpreting the model parameters from constants, ie. ‘literals’, to model states with derivative 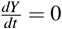. Any parameter that needs calibration so that the model displays a certain value for a variable (target value) has its derivative function 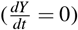 replaced with any another monotonic function that would respect the correlations of table 4. In this work, parameter adaptation is implemented using a cubic correction law

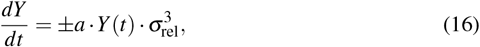

where *a* is a scaling factor and *σ*_rel_ = (*Y*_target_ − *Y*_current_)*/Y*_target_ represents the relative error between the current model output and its target value. The sign of the correction term is selected to ensure consistency with the qualitative parameter/variable relationships.

Inspecting the equation we observe that if the relative error is high, the value used for the parameter derivative is very high and as the error reaches 0, the value for the derivative will also be 0. Here an error of 0 means that the model is displaying the same value for the variable being calibrated as the target value for that variable. To maintain the system within operational bounds, a *Y*_*max*_ and *Y*_*min*_ are defined for the parameter. The maximum value for the error is also capped at ± 1 to cap the maximum value of the derivative. To rescale the error value to the dimensions of the parameter, the current value *Y* of the parameter is used. The coefficient *a* acts as a gain parameter that scales the responsiveness of the calibration dynamics, controlling the speed and aggressiveness of parameter adaptation.

#### 2.3.2 Construction of the full calibration system

We now extend the Embedded Gradient Descent (EGD) strategy from the Generic Compartment to the full cardiopulmonary circulation. Using the model of Figure 2 as a reference, our goal is to construct a complete set of controller equations that can calibrate the system to any admissible set of measured variables. The selection of these controllers is guided by three elements: the structure of the governing equations, the qualitative parameter/variable correlations summarized in Table 4, and basic physiological principles. The resulting mapping between targets and parameters is shown in Figure 6, organized into ten calibration groups.

**Figure 6.**
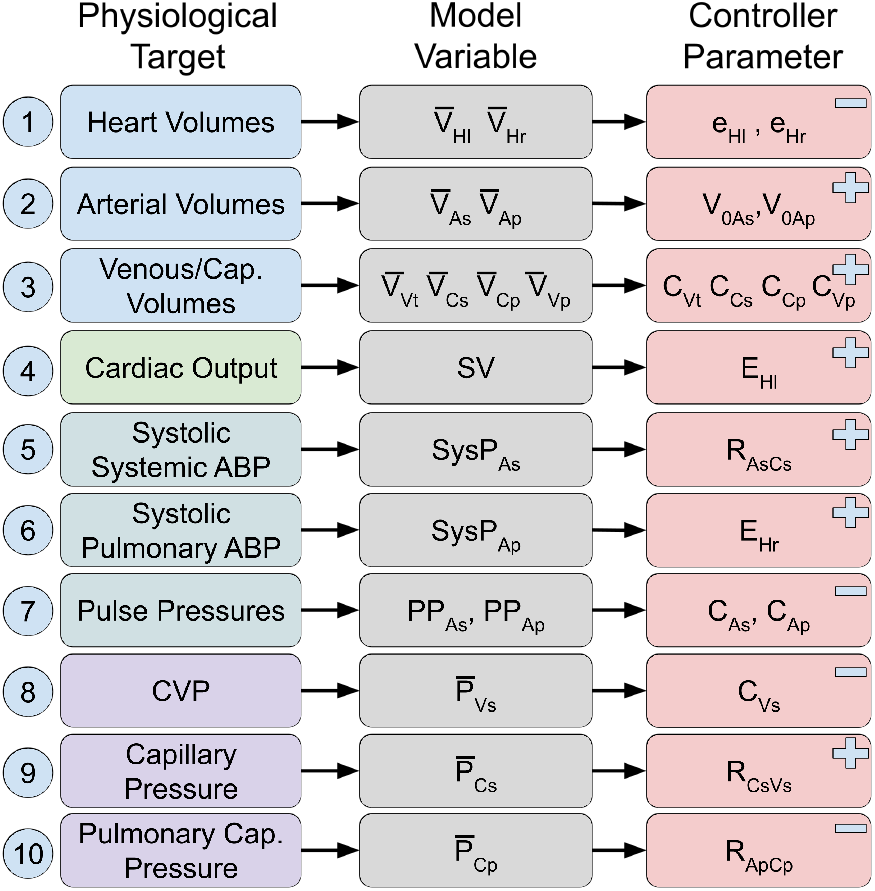
Overview of the parameter/target assignments used in the Embedded Gradient Descent calibration. Each row corresponds to a physiological quantity targeted during calibration (left), the associated model variable (centre) and the parameter adjusted by the controller (right). Symbols ‘+’ and ‘−’ denote positive and negative qualitative correlations between parameters and target variables, respectively, as defined in Table 4.

#### Volumes equations (groups 1 to 3)

Because TBV is fixed, only eight of the nine compartmental volumes listed in Table 1 must be actively targeted. Therefore, the systemic venous compartment is left uncontrolled so that it absorbs the residual volume required to satisfy conservation. For the heart chambers (group 1), the average diastolic volumes are controlled using the relaxed elastances *e*_*Hl*_ and *e*_*Hr*_: among the cardiac parameters, these exert the strongest and most selective influence on diastolic filling, with a clear negative correlation between elastance and volume. The arterial volumes (group 2) are regulated through their unstressed volumes *V*_0,*As*_ and *V*_0,*Ap*_, the only compartments for which *V*_0_ ≠ 0. Because *V*_0_ shifts the pressure–volume relation in Eq. 1 without altering compliance, it provides a direct and isolated mechanism for adjusting the mean arterial volume. Venous and capillary volumes (group 3) are controlled through their compliances (*C*_*Vt*_, *C*_*Cs*_, *C*_*Cp*_, *C*_*V p*_), which appear in Eq. 1 and exert a positive, dominant influence on stored volume in these highly compliant regions of the circulation. These parameter choices ensure that each targeted volume is adjusted through the variable with the most physiologically direct and least confounded effect on that compartment.

#### Cardiac output equation (group 4)

Stroke volume, and thus cardiac output, is controlled through the left ventricular contraction elastance *E*_*Hl*_. Because this parameter determines the amplitude of ventricular emptying, it provides a direct way to reach the target stroke volume while remaining consistent with the overall assumption of uniform average flow throughout the system.

#### Arterial pressures and pulse pressures (groups 5 to 7)

Systemic systolic pressure (group 5) is regulated with the systemic outlet resistance *R*_*AsCs*_, as the inlet resistance is assumed small and fixed. Once systolic pressure is set, the corresponding pulse pressure (group 7) is shaped by the systemic arterial compliance *C*_*As*_, whose negative correlation with pressure amplitude makes it well suited for this role. The same logic is applied symmetrically in the pulmonary circulation: the right ventricular contraction elastance *E*_*Hr*_ determines pulmonary systolic pressure (group 6), while pulmonary arterial compliance *C*_*Ap*_ is used to adjust the pulmonary pulse pressure (group 7).

#### Venous and capillary pressures (groups 8 to 10)

Central venous pressure (group 8) is controlled through the systemic venous compliance *C*_*Vs*_, which modulates the pressure/volume relation in this compartment. Systemic capillary pressure (group 9) is influenced most strongly by the downstream resistance *R*_*CsVs*_ and is therefore calibrated using this parameter. In the pulmonary capillaries (group 10), the inlet resistance *R*_*ApCp*_ typically dominates the local pressure drop, making it the appropriate choice for controlling *P*_*Cp*_.

#### Combined effect of all controllers

This assignment yields a set of sixteen controller equations corresponding to the sixteen extended targets in Figure 6. Each target is paired with the parameter that most directly and selectively influences it, following the qualitative correlations of Table 4. Two regions of the circulation, the thoracic veins (*P*_*Vt*_) and pulmonary veins (*P*_*V p*_), are intentionally left uncontrolled. Although resistances connect these compartments to their neighbours, these venous resistors, together with all cardiac resistors, are assumed to be very small, producing negligible pressure drops and modifying them would therefore have minimal physiological effect. When the full set of controllers is active, all sixteen equations operate simultaneously, with each correction altering pressures and flows throughout the network. The resulting interactions cause the controllers to continually adjust one another, guiding the system toward the configuration in which all relative errors are minimized.

#### 2.3.3. Construction of calibration targets from model outputs

Clinical observables and derived haemodynamic targets (e.g., systolic pressure, pulse pressure, stroke volume, and mean compartmental volumes) are defined over cardiac cycles rather than at instantaneous time points. To enable direct comparison between continuous model states and these cycle-resolved targets, the auxiliary operators introduced in Section 2.2.3 are designed to extract clinically meaningful quantities online during numerical integration.

Stroke volume is computed as the value of the cycle-integrated aortic flow retained from the previous heartbeat,

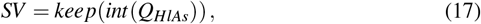

and cardiac output is then obtained from *SV* and heart rate as *CO* = *SV* · *HR* (with consistent unit conversion where required). Systolic arterial pressure is extracted as the retained cycle maximum,

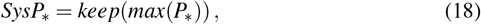

while pulse pressure is computed from the retained cycle extrema,

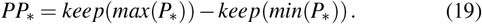

Compartmental volume targets are formulated using cycle-averaged values,

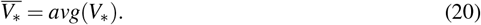

The complete set of operator compositions used to construct the calibration targets, together with the corresponding model variables and the parameters adapted by each controller, is summarised in Fig. 6. This mapping defines the interface between clinical targets and the internal model dynamics within the Embedded Gradient Descent framework.

### 2.4 Numerical implementation details

All differential equations defining the cardiovascular model, cycle-based operators, and calibration dynamics were integrated using an explicit Euler scheme with a fixed time step Δ*t*. This formulation was selected to allow precise control over event-like behaviour, such as cardiac cycle triggering and operator resets, which are naturally expressed within a time-stepped framework.

Discrete events, including cardiac cycle triggers and the reinitialisation of cycle-based operators (e.g., maxima, minima, and integrals), were implemented as fast relaxations over a single integration step rather than as explicit hybrid events. This approach avoids discontinuities in the state variables while preserving the intended cycle-level behaviour within a purely ODE-based formulation.

The integration time step was set to Δ*t* = 0.0005 s, which was sufficient to resolve the fastest dynamics in the system, including elastance transitions and calibration updates, while maintaining computational efficiency and numerical stability.

Parameter bounds were introduced solely as numerical safeguards to prevent divergence. These bounds were intentionally chosen to be substantially wider than physiologically plausible ranges and therefore do not constrain the solution space in a meaningful way. Physiological plausibility emerges from the governing equations and their coupling structure rather than from imposed parameter limits.

### 2.5 Simulation Setup

To assess whether the calibration framework converges to the same solution independently of initial conditions, 100 simulations were performed in which model parameters were initialised randomly and TBV was distributed randomly across compartments. All simulations were run for 2200 s of simulated time, allowing the coupled physiological and calibration dynamics to reach a steady regime.

For the convergence analysis, target values corresponded to the average normal-physiology subject defined in Table 5. Each simulation included an initial 200 s transient phase without active controllers, allowing the system to settle and preventing large initial parameter excursions. The remaining 2000 s were simulated with controllers enabled, which was found sufficient for both state variables and calibrated parameters to converge.

**Table 5.**
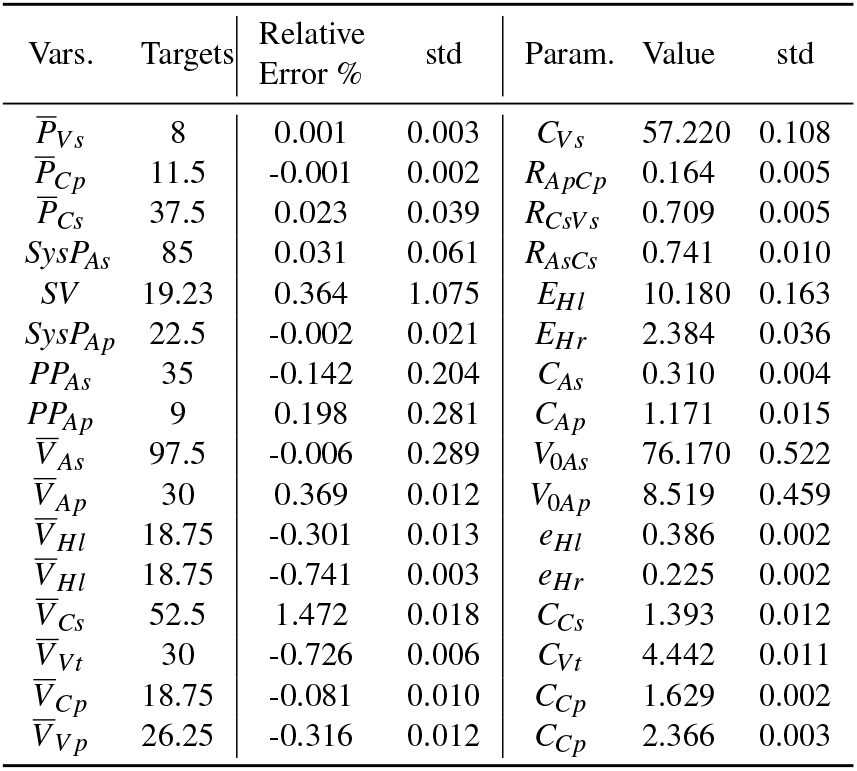
Summary of the convergence test across 100 simulations with random initial parameter values and volume distributions. For each calibration target, the table reports the prescribed target value, the mean signed relative error and its standard deviation at convergence, together with the corresponding calibrated parameter values and their variability across runs.

| Vars. | Targets | Relative Error % | std | Param. | Value | std |
| --- | --- | --- | --- | --- | --- | --- |
| $\bar{P}_{Vs}$ | 8 | 0.001 | 0.003 | $C_{Vs}$ | 57.220 | 0.108 |
| $\bar{P}_{Cp}$ | 11.5 | -0.001 | 0.002 | $R_{ApCp}$ | 0.164 | 0.005 |
| $\bar{P}_{Cs}$ | 37.5 | 0.023 | 0.039 | $R_{CsVs}$ | 0.709 | 0.005 |
| $SysP_{As}$ | 85 | 0.031 | 0.061 | $R_{AsCs}$ | 0.741 | 0.010 |
| $SV$ | 19.23 | 0.364 | 1.075 | $E_{Hl}$ | 10.180 | 0.163 |
| $SysP_{Ap}$ | 22.5 | -0.002 | 0.021 | $E_{Hr}$ | 2.384 | 0.036 |
| $PP_{As}$ | 35 | -0.142 | 0.204 | $C_{As}$ | 0.310 | 0.004 |
| $PP_{Ap}$ | 9 | 0.198 | 0.281 | $C_{Ap}$ | 1.171 | 0.015 |
| $\bar{V}_{As}$ | 97.5 | -0.006 | 0.289 | $V_{0As}$ | 76.170 | 0.522 |
| $\bar{V}_{Ap}$ | 30 | 0.369 | 0.012 | $V_{0Ap}$ | 8.519 | 0.459 |
| $\bar{V}_{Hl}$ | 18.75 | -0.301 | 0.013 | $e_{Hl}$ | 0.386 | 0.002 |
| $\bar{V}_{Hr}$ | 18.75 | -0.741 | 0.003 | $e_{Hr}$ | 0.225 | 0.002 |
| $\bar{V}_{Cs}$ | 52.5 | 1.472 | 0.018 | $C_{Cs}$ | 1.393 | 0.012 |
| $\bar{V}_{Vt}$ | 30 | -0.726 | 0.006 | $C_{Vt}$ | 4.442 | 0.011 |
| $\bar{V}_{Cp}$ | 18.75 | -0.081 | 0.010 | $C_{Cp}$ | 1.629 | 0.002 |
| $\bar{V}_{Vp}$ | 26.25 | -0.316 | 0.012 | $C_{Cp}$ | 2.366 | 0.003 |

To evaluate the ability of the framework to generate heterogeneous yet physiologically consistent populations, three cohorts were constructed corresponding to normal physiology, warm shock (WS), and cold shock (CS). For each cohort, admissible target ranges were defined according to Table 2. A Latin hypercube sampling strategy was used to generate 200 independent target sets per cohort. Each target set was then used to calibrate the full cardiovascular model using the Embedded Gradient Descent procedure, yielding one virtual subject per sample. The same simulation protocol and timing configuration were used for all cohorts.

## 3 RESULTS

### 3.1 Convergence analysis

Table 5 summarises the results of the calibration procedure. Across all simulations, all targeted variables converged to their prescribed values with negligible signed relative error (below 1%, computed as 100 × (*Y*_value_ − *Y*_target_)*/Y*_target_). The corresponding distributions of absolute calibration errors are shown in Fig. 7 for each calibrated variable. For all targets, the absolute error at convergence remained within ± 0.75 mmHg for pressure and volume variables.

**Figure 7.**
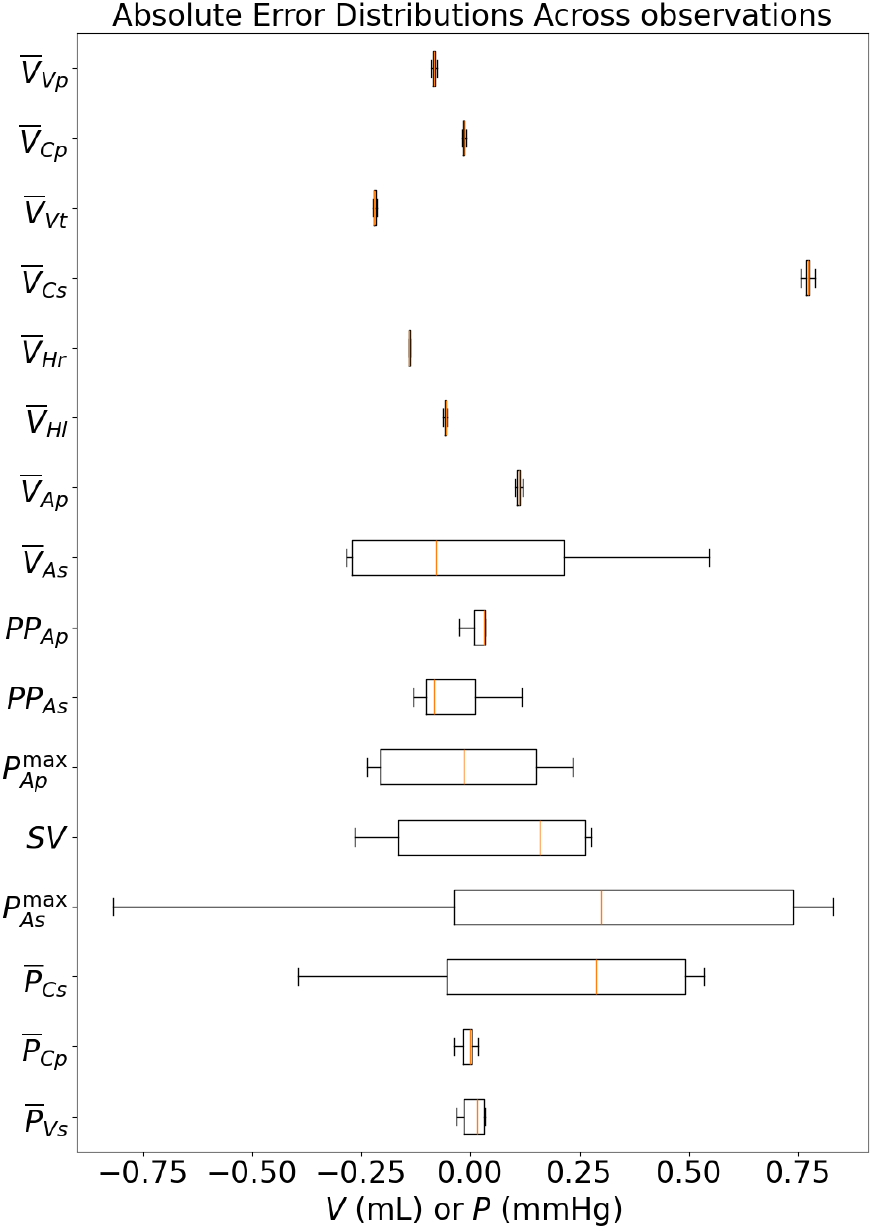
Distribution of absolute calibration errors across all targeted physiological variables at convergence. Errors are reported in mmHg for pressure targets, mL · for volume targets and *mL · beat*^−1^ for stroke volume. All simulations converge to a narrow error band despite random initialization of parameters and compartmental volume distributions.

### 3.2 Sepsis Population

To evaluate the ability of the calibration framework to generate heterogeneous yet physiologically consistent populations, three cohorts were constructed corresponding to normal physiology, warm shock (WS), and cold shock (CS). For each cohort, admissible target ranges were defined according to Table 2. A Latin hypercube sampling strategy was used to generate 200 independent target sets per cohort. Each target set was then used to calibrate the full cardiovascular model using the Embedded Gradient Descent procedure, yielding one virtual subject per sample.

Figure 8 summarises the distribution of relative calibration errors across all targeted pressures, volumes, and cardiac output for the three cohorts. For the majority of variables, errors are tightly centred around zero and symmetrically distributed, showing that the calibration procedure reliably achieves the prescribed targets across a wide range of physiological conditions.

**Figure 8.**
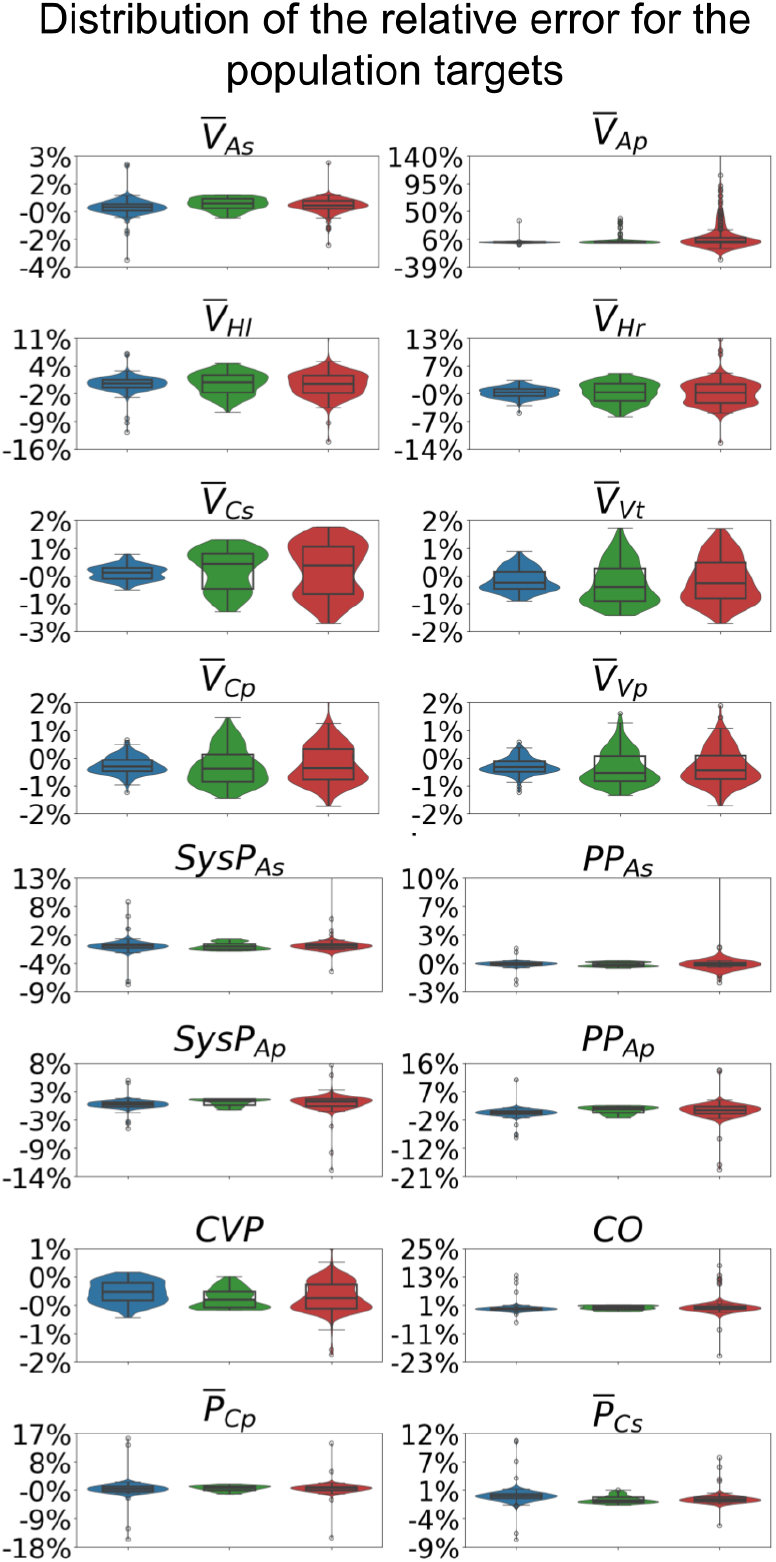
Distribution of relative calibration errors across all targeted physiological variables for the normal, warm shock (WS), and cold shock (CS) populations. Errors are computed as relative deviations from the prescribed targets after convergence. Most variables exhibit errors tightly centred around zero, indicating robust calibration across cohorts.

The first pair of plots in Fig. 9 focuses on the pulmonary arterial volume 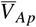, which exhibits the largest residual errors across the population. These errors are observed predominantly in the warm shock cohort. Importantly, the error is consistently positive, indicating a systematic tendency towards increased pulmonary arterial volume relative to the prescribed target. As shown in the corresponding parameter space, this behaviour coincides with saturation of the unstressed pulmonary arterial volume parameter *V*_0,*Ap*_ at its imposed lower bound. The second pair of plots highlights a similar saturation phenomenon affecting cardiac controllers. In this case, large residual errors are again associated with parameter saturation, but now lead predominantly to negative errors. In contrast, the third pair of plots shows a regime in which no controller saturation is observed and the error distribution is smoothly spread across the population.

**Figure 9.**
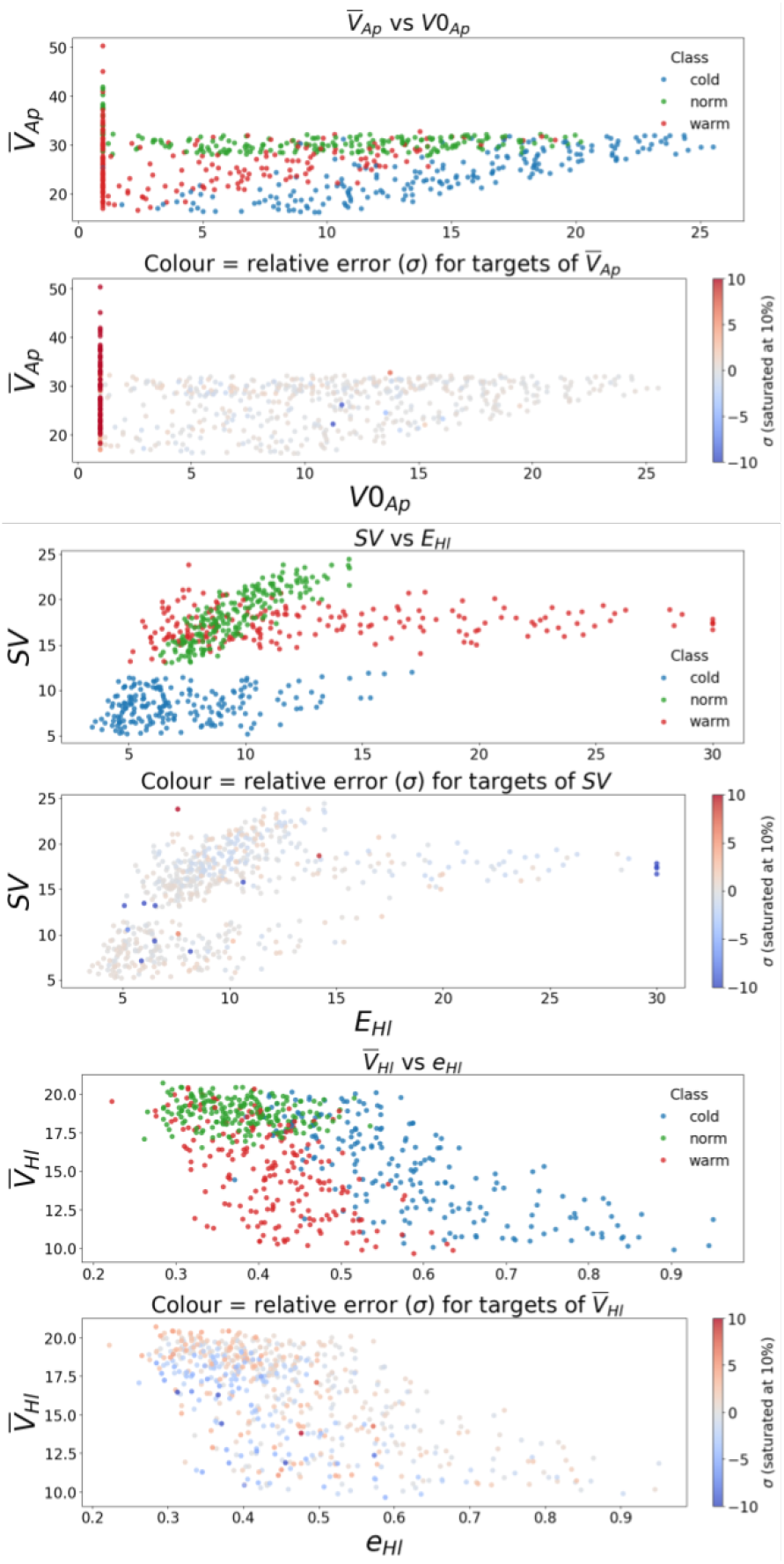
Relative calibration errors for representative population parameters across normal, warm shock, and cold shock cohorts. Upper panels show parameter values coloured by class; lower panels show the same samples coloured by relative error (saturated at 10%). Large residual errors coincide with controller saturation in specific regimes, while non-saturating controllers exhibit smoothly distributed errors driven by the cubic control law.

### 3.3 Population Parameters

Figure 10 shows how global vascular and cardiac parameters organise across the three cohorts as functions of TBV and cardiac output. The top two panels show the distribution of total systemic resistance as a function of TBV and cardiac output. Across all cohorts, *R*_*Total*_ exhibits a strong negative correlation with cardiac output and only a weak dependence on total blood volume. Despite this shared flow dependence, the cold shock population consistently occupies higher resistance regimes than normal physiology, while the warm shock population exhibits systematically lower resistance at comparable cardiac output and blood volume.

**Figure 10.**
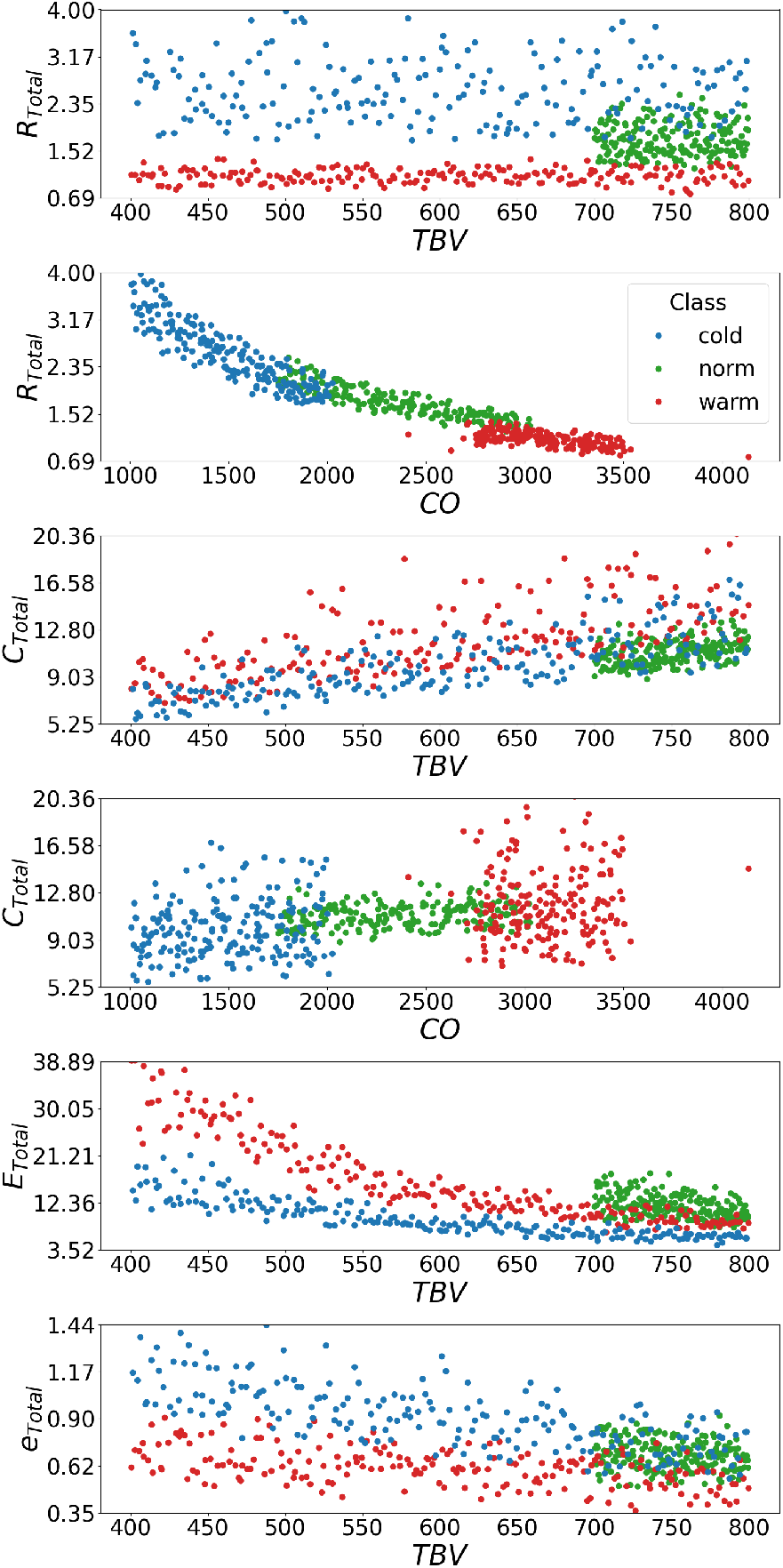
Distribution of total systemic resistance (*R*_*Total*_), compliance (*C*_*Total*_), and cardiac elastance (*E*_*Total*_ and *e*_*Total*_) across normal, warm shock, and cold shock populations as a function of total blood volume (*TBV*) and cardiac output (*CO*).

The third and fourth panels illustrate the total system compliance also as a function of TBV and cardiac output. Across all cohorts, *C*_*Total*_ increases with total blood volume, whereas no clear dependence between *C*_*Total*_ and cardiac output is observed. On average, the warm shock population occupies higher compliance regimes than the cold shock population, and for configurations with normal blood volumes, both septic populations show higher compliance than the normal cohort. Within the cold shock population at normal blood volumes, the total systolic elastance *E*_*Total*_ (5th panel) is markedly reduced, while the total diastolic elastance *e*_*Total*_ (6th panel) remains comparable to that of the normal population. Across the full volume range, *e*_*Total*_ in cold shock is consistently higher than in warm shock, and in the low-volume regime both septic populations exhibit elevated diastolic elastance relative to normal physiology. Conversely, in the low-volume regime, the warm shock population exhibits substantially elevated systolic elastance relative to normal physiology.

Figure 11 illustrates the relationship between the arterial/capillary resistance ratio,

**Figure 11.**
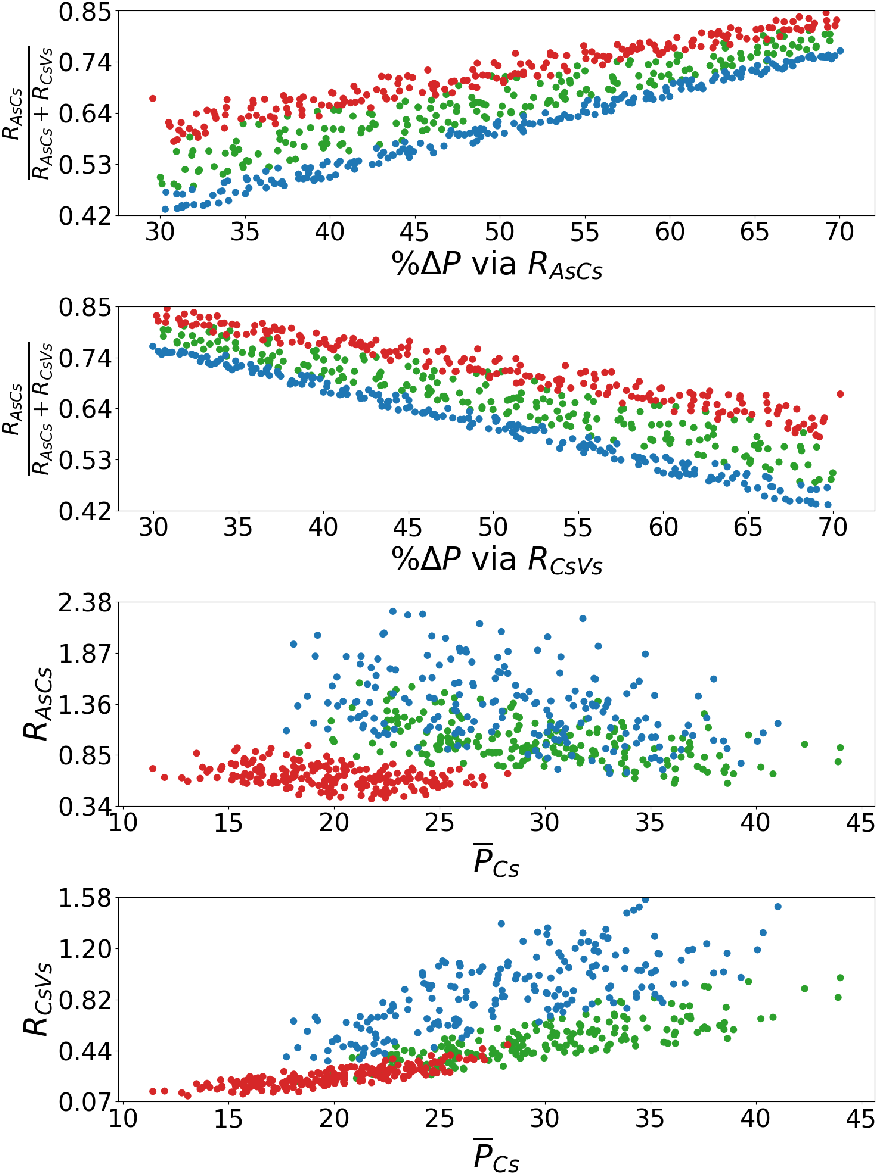
Top: Relationship between the arterial–capillary resistance ratio and the fractional pressure drop across arterial and venous compartments for normal, warm shock, and cold shock populations. Bottom: Scatter plots between the arterial resistance (*R*_*AsCs*_) and the venous resistance (*R*_*CsVs*_) and the average pressure of the capillaries 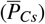.

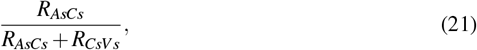

and the fractional pressure drop occurring across the arterial segment,

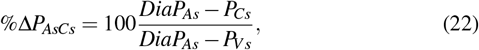

as well as across the venous segment,

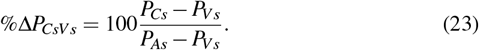

The top panel shows that higher values of the resistance ratio correspond to a greater fraction of the pressure drop occurring between the arteries and the capillaries. Conversely, the second panel demonstrates complementary behaviour on the venous side, where increasing resistance ratios lead to a reduced fraction of the total pressure drop occurring between the capillaries and the veins. In the present results, the resistance ratio for the warm shock population is systematically higher than that of the cold shock population for comparable fractional pressure drops. The third and fourth panels relate the individual inlet and outlet resistances to the mean capillary pressure. In the third panel, *R*_*AsCs*_ shows only a weak association with 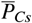: the warm shock population occupies a narrow, low-*R*_*AsCs*_ band at lower 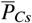, whereas cold shock spans a much wider resistance range with a tendency for higher 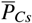 to coincide with lower *R*_*AsCs*_.

In contrast, the fourth panel shows a clear monotonic increase of venous resistance *R*_*CsVs*_ with 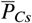 across all cohorts, with the largest values occurring in cold shock. Together, these panels indicate that the effective resistance partitioning (and therefore the resistance ratio) is driven primarily by variations in *R*_*CsVs*_, while *R*_*AsCs*_ remains comparatively constrained and does not provide a consistent monotonic signature with capillary pressure.

In addition to reproducing the targeted haemodynamic states, the generated populations were assessed for consistency with the qualitative correlation structure assumed during model formulation (Table 4). In Fig. 10, the total systemic resistance shows no clear dependence on TBV and exhibits a strong negative correlation with cardiac output. The same figure shows that total compliance increases with blood volume, whereas total elastance decreases.

Figure 11 shows that the ratio *R*_*AsCs*_*/*(*R*_*AsCs*_ + *R*_*CsVs*_) increases with the fraction of pressure dropped across the arterial segment and decreases with the fraction dropped across the venous segment. The lower panels show the relationships between *R*_*AsCs*_, *R*_*CsVs*_ and 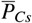 in the calibrated populations.

Finally, Fig. 12 shows that capillary compliance (*C*_*Cs*_) decreases with increasing mean capillary pressure 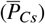, arterial compliance (*CAs*) decreases with pulse pressure (*PP*_*As*_), and unstressed arterial volume (*V*_0,*As*_) increases with mean arterial volume 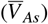. The relationship between stroke volume and left-heart systolic elastance (*E*_*Hl*_) is also shown.

**Figure 12.**
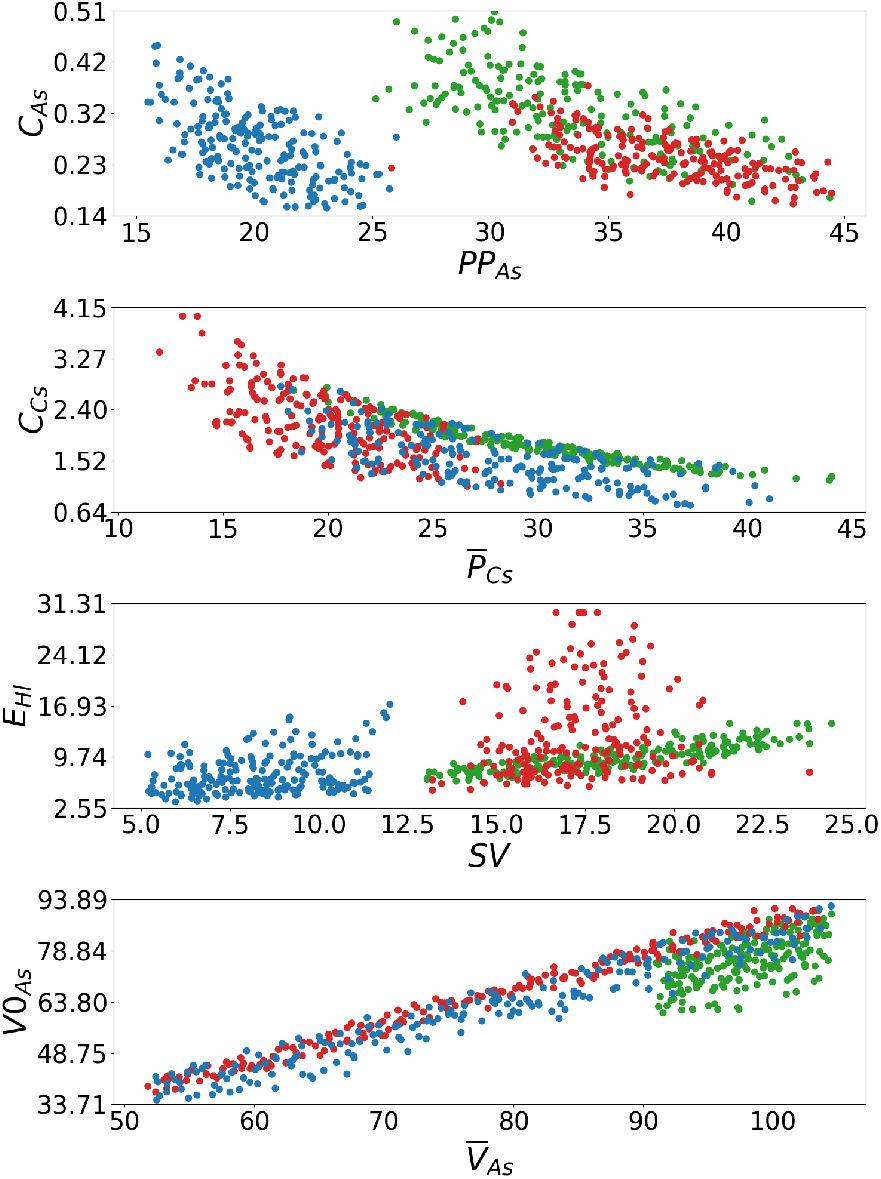
Scatter plots between arterial systemic compliance (*C*_*As*_) and arterial pulse pressure (*PP*_*As*_), systemic capillary compliance (*C*_*Cs*_) and average capillary pressure 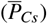 systolic elastance of the left ventricle (*E*_*Hl*_), and Stroke Volume (*SV*) and unstressed volume of the Systemic artery (*V*_0*As*_) and average volume of the systemic arteries 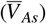.

## 4 DISCUSSION

### Convergence analysis

The small residual errors observed at convergence reflect the fact that calibration is performed on a fully dynamic, cycle-resolved system. Targets such as systolic pressure, pulse pressure, stroke volume, and cycle-averaged volumes are computed using max/min/average operators over the cardiac cycle, rather than representing static state variables. As a consequence, controlled variables and their associated parameters do not converge to static values but instead exhibit small, bounded oscillations around the target equilibrium, as illustrated in Fig. 13. These oscillations arise from the cycle-resolved nature of the calibration and remain well within negligible ranges. A representative calibration trajectory across multiple time scales is provided in Appendix A.3. Taken together, these findings indicate that the Embedded Gradient Descent dynamics are robust to initial parametrisation and volume partitioning, converging reproducibly to a unique steady solution for a given target set.

**Figure 13.**
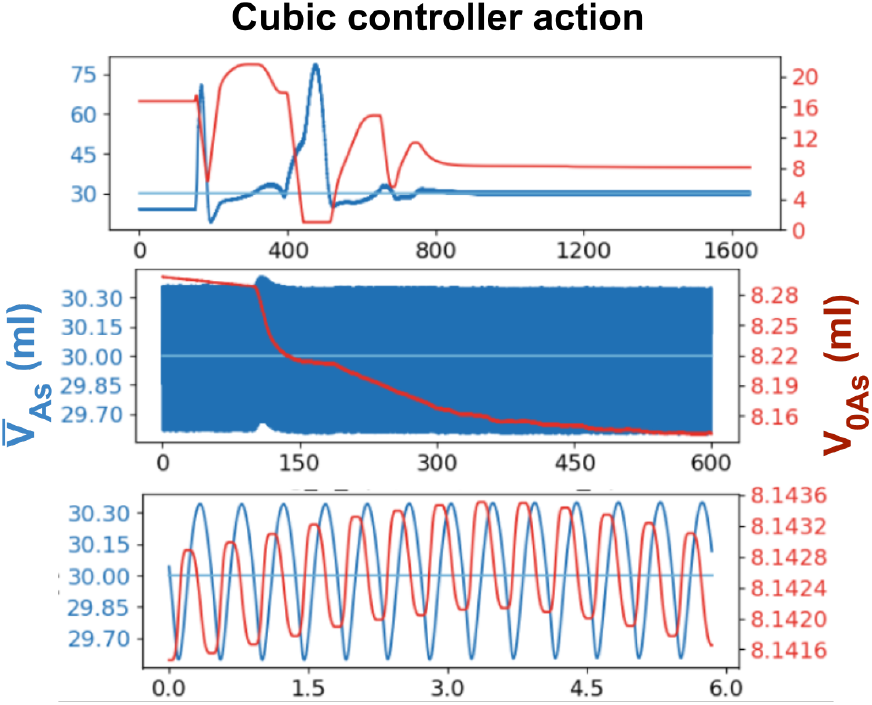
Calibration trajectory for the average systemic arterial volume 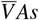 (blue) and the corresponding unstressed volume parameter *V* 0, *As* (red), shown across three time scales. Top: full calibration run, illustrating large transient parameter adjustments during the initial convergence phase. Middle: intermediate time window highlighting the gradual approach of 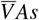 towards its prescribed target and the associated monotonic adaptation of *V* 0, *As*. Bottom: short time-scale view at convergence, showing bounded, cycle-resolved oscillations of both variables around the target equilibrium. These oscillations arise from the fully dynamic nature of the system and remain within clinically negligible ranges.

### Solution uniqueness

The convergence behaviour demonstrated in the previous section is a consequence of a deeper structural property of the calibration system, namely the uniqueness of the physiological solution enforced by the Embedded Gradient Descent framework.

A key property of the EGD calibration procedure is that, for any admissible set of targets, it converges to a single physiological solution. Although 16 parameters are calibrated, each is tied to a specific target through the correlations of Table 4, and each controller acts as a monotonic attractor that pulls its assigned variable toward the desired value. Because every parameter influences multiple compartments, each controller imposes a global constraint on the circulation and the system must also simultaneously satisfy the intrinsic mathematical relations of the ODE model. The resulting problem is strongly over-determined: most parameter combinations violate one or more constraints, and the only possible equilibrium is the configuration at which all relative errors *σ*_rel_ vanish. With all attractors active at once, any perturbation of one parameter immediately alters the conditions seen by the others, preventing convergence to partial or local minima.

Figure 14 provides an illustrative low-dimensional example of the constraint structure induced by the EGD calibration. In this representation, pressure is plotted as a function of arterial compliance and volume according to Eq. 1. When compliance is used to regulate pressure in a given compartment, the corresponding controller defines a one-dimensional set of admissible parameter states for which the pressure target is satisfied (*σ*_rel_ = 0, black curve), independent of the remainder of the system. Simultaneously, constraints imposed by the other controllers restrict the compartment to evolve along an independent trajectory (yellow curve). The calibrated solution corresponds to the unique state satisfying both constraints, indicated by their intersection (red marker).

**Figure 14.**
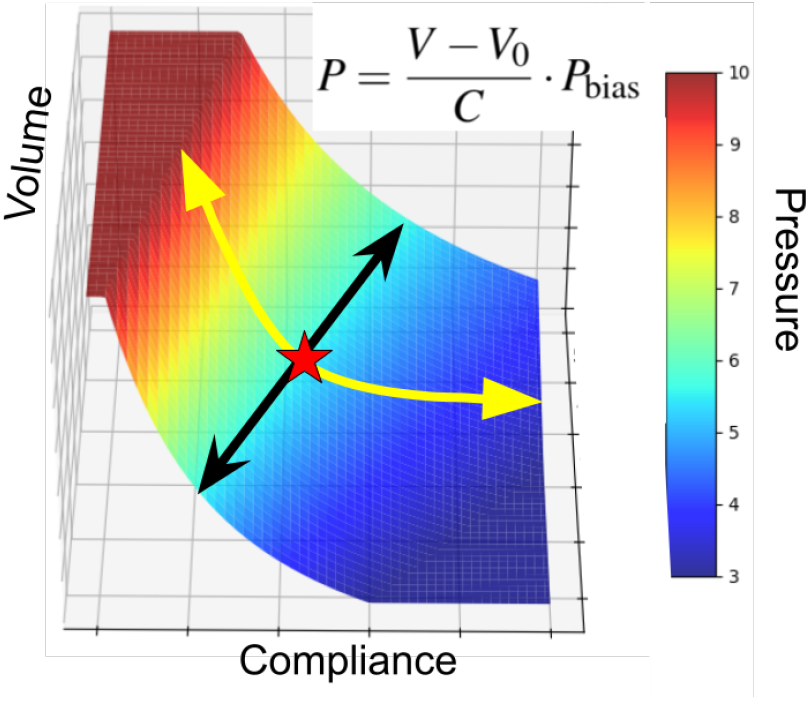
Conceptual illustration of solution uniqueness in the Embedded Gradient Descent calibration framework. Pressure is shown as a function of compartmental volume and compliance according to Eq. 1. One controller enforces the pressure target, defining a set of admissible parameter combinations (black curve), while constraints imposed by the remaining controllers restrict the system to an independent trajectory (yellow curve). The calibrated solution corresponds to the unique state satisfying both constraints simultaneously (red marker).

In the full cardiovascular model, the same principle applies in a higher-dimensional parameter space: each controller restricts the system to a subset of admissible states, and the simultaneous action of all controllers enforces convergence to their unique common intersection.

If a particular target set is incompatible with the model, one or more controllers drive their parameters to their bounds *y*_min_ or *y*_max_ while the corresponding *σ*_rel_ remains non-zero. A saturated controller behaves as a constant parameter, effectively removing one attractor from the system. The remaining controllers still minimise their errors subject to this constraint, yielding a best-compromise equilibrium in which feasible targets are satisfied and infeasible ones are highlighted by persistent residual errors and parameters pinned at their limits.

### Sepsis Population

The error patterns observed in Fig. 9 provide insight into how the calibration framework responds when prescribed targets become incompatible with the governing equation system. In the first pair of plots, saturation of the unstressed pulmonary arterial volume parameter *V*_0,*Ap*_ prevents further reduction once its lower bound is reached. At this point, the corresponding controller loses authority, rendering the target for 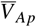 unattainable in this regime. The calibration therefore proceeds with one fewer active constraint, while the remaining controllers continue to minimise their respective errors. The resulting persistent residual error is not indicative of algorithmic failure, but instead serves as a direct diagnostic of structural incompatibility between the imposed target combination and the model formulation.

A comparable mechanism is evident in the second pair of plots, where large negative residuals are associated with saturation of parameters governing cardiac contractility. In this case, the reduced control authority limits the model’s ability to increase stroke volume under the prescribed targets. Importantly, these saturation effects remain confined to specific regions of the parameter space, while the majority of the population exhibits small, unbiased errors. This indicates that the observed residuals reflect local feasibility limits rather than a loss of global calibration performance.

In contrast, the third pair of plots in Fig. 9 illustrates a regime in which no controller saturation occurs and residual errors are smoothly distributed across the population. Here, convergence behaviour is dominated by the cubic control law, which produces progressively weaker corrective action as the error magnitude decreases. This leads to slow convergence near the target and the persistence of small residual errors of either sign. The direction of the final residual depends on the initial conditions, with trajectories approaching the target from either side.

Taken together, these observations show that large residual errors arise only when target specifications exceed the structural limits of the model, which are made explicit through parameter saturation. Outside these regimes, the calibration framework yields well-behaved error distributions while transparently exposing the boundaries of physiological feasibility imposed by the model structure.

### Population Parameters

The WS cohort exhibits low resistance across the explored volume range (Fig. 10), consistent with near maximal systemic vasodilation, whereas the cold shock cohort spans higher and more variable resistance states, consistent with vasoconstrictive responses (Fig. 1). Although warm shock also exhibits, on average, higher total compliance than cold shock, both septic cohorts show elevated compliance relative to normal physiology in the normal volume range, which is expected for WS but not for CS. The elastance panels suggest an additional cardiac contribution: at normal blood volumes, cold shock shows a marked reduction in total systolic elastance *E*_*Total*_ relative to normal, while diastolic elastance *e*_*Total*_ remains broadly comparable, a pattern consistent with impaired contractile strength with preserved diastolic properties and therefore compatible with a cardiogenic component. Conversely, in the low-volume regime the warm shock cohort exhibits substantially elevated *E*_*Total*_, implying a configuration that would require disproportionately high systolic stiffness to sustain flow under combined hypovolaemia and vasodilation, and is therefore unlikely to represent a stable haemodynamic state over time.

From a physiological perspective, warm shock is characterised by a state of generalized vasodilation, which primarily affects the arterial compartment, while venous resistance is typically less responsive. Under such conditions, one would expect a lower arterial/capillary resistance ratio and a larger fraction of the total pressure drop to occur downstream, at the venous level. In contrast, cold shock is associated with arterial vasoconstriction, leading to higher resistance ratios and a larger pressure drop upstream. However, inspection of the parameter distributions of figure 11 indicates that this behaviour arises primarily from variations in venous resistance, while arterial resistance in WS remains comparatively constrained across populations. As a result, changes in the resistance ratio are driven predominantly by the venous compartment rather than by arterial vasodilation. This parameter distribution reflects the way population targets were specified. Identical fractional pressure-drop ranges (30–70%) were imposed for both warm and cold shock cohorts, while systolic arterial pressure targets were also shared across populations. At the same time, higher pulse pressure targets in warm shock imply lower diastolic arterial pressures, which, combined with percentage-based capillary pressure targets relative to diastolic pressure and CVP, systematically bias the warm shock population towards lower capillary pressure targets. A more physiologically faithful separation between warm and cold shock could be achieved by assigning lower arterial pressure-drop ranges to the warm shock population and higher ranges to the cold shock population, thereby explicitly biasing the resistance ratio towards arterial vasodilation or vasoconstriction, respectively.

Beyond matching the target haemodynamic ranges, the calibrated populations broadly preserve the qualitative correlation structure assumed during model formulation (Table 4), which provides an internal consistency check on the generated parameter space. In Fig. 10, the inverse coupling between *R*_*Total*_ and cardiac output is maintained across cohorts, while the increase of *C*_*Total*_ with TBV and the concomitant reduction in effective elastance with filling reflect the expected behaviour of compliant compartments under volume loading. Figure 11 further demonstrates that the resistance partitioning remains mechanically coherent: the ratio *R*_*AsCs*_*/*(*R*_*AsCs*_ + *R*_*CsVs*_) varies monotonically with the prescribed fractional pressure-drop allocations, consistent with resistors arranged in series. Also in agreement with the correlation assumptions, Fig. 11 shows that the arterial resistance *R*_*AsCs*_ (inlet resistor) is positively correlated with the mean capillary pressure 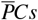, while the venous resistance *RCsVs* (outlet resistor) is negatively correlated with the mean capillary pressure 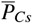. Finally, Fig. 12 shows that the imposed local compliance/pressure and compliance/pulse-pressure relationships are retained (*C*_*Cs*_ decreasing with 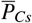 and *C*_*As*_ decreasing with *PP*_*As*_), and that the scaling between *V*_0,*As*_ and 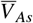 remains approximately linear across the sampled space. Taken together, these patterns support that the calibrated populations are not arbitrary collections of parameters but occupy a physiologically structured manifold consistent with the model’s embedded assumptions.

### Limitations

Despite the strengths of the proposed approach, several limitations should be acknowledged. First, calibration was performed for steady-state snapshots rather than continuous time series, such that transient dynamics were not explicitly represented, for example progressive total blood volume loss due to vascular leakage. While this choice enabled systematic population-level analysis and robust convergence assessment, it limits direct inference of short-term physiological adaptations. Future work will therefore focus on applying the method to real patient recordings and on analysing the temporal evolution of inferred physiological parameters in response to disease progression and therapeutic interventions. In this context, extending the model beyond purely haemodynamic variables will be essential. Sepsis is characterised not only by circulatory dysfunction but also by metabolic derangements, such as elevated lactate, and impaired gas exchange, for which the capillary network plays a central physiological role. Incorporating blood gas fractions, lactate kinetics, and membrane transport and equilibrium dynamics would provide additional constraints on capillary behaviour and help inform the structural and functional characteristics required to represent microcirculatory function more faithfully. Second, population generation assumed a common reference distribution of total blood volume across compartments. While this choice supports direct comparability of inferred parameter sets across individuals and physiological states, the results indicate that it does not eliminate the need to explore controlled deviations from the normal distribution. In particular, persistent residual errors in arterial volume–related targets, such as *V*_*Ap*_, highlight regimes in which the imposed volume partitioning is not fully compatible with the governing equation system. Under pathological conditions such as hypovolaemia and septic shock, physiological adaptation is expected to involve redistribution of blood volume between vascular compartments, making systematic variation of volume partitioning a necessary extension of the present framework. Lastly, the present model does not include inertial (inductive) effects of blood flow; while this simplification has negligible impact on steady-state calibration, such effects become relevant under dynamic conditions, where frequency-dependent responses and resonance phenomena may influence transient haemodynamics and should be considered in future extensions.

### Conclusion

In this work, we developed and validated a mechanistic cardiovascular model coupled to an Embedded Gradient Descent (EGD) calibration framework capable of generating physiologically consistent solutions across multiple haemodynamic states, including normal physiology as well as warm and cold septic shock. The proposed calibration strategy consistently drives the model towards the prescribed physiological targets and yields unique calibrated solutions with low residual error across a broad range of physiological conditions. The use of a common blood volume distribution across all populations was a deliberate design choice, ensuring that calibrated parameter sets remain directly comparable between individuals and across physiological states. This uniformity is essential for population-level analyses and for attributing observed differences to underlying haemodynamic adaptations.

A further important outcome of this work is that residual calibration errors provide interpretable information about the internal consistency of the model/target configuration. In particular, persistent residuals occur only when specific target combinations are incompatible with the governing constraints, and are directly associated with controller saturation, thereby exposing feasibility limits in an explicit and interpretable manner. Interpreting residual error as a diagnostic of structural feasibility, rather than as optimisation failure, directly reflects the methodological principles advocated in recent guidelines for mechanistic cardiovascular modelling, which emphasise physiological interpretability, explicit handling of identifiability limits, and the integration of calibration within the governing model structure rather than as an external black-box procedure [27].

An additional advantage of the EGD-based calibration approach is its scalability. Unlike traditional optimisation-based calibration methods, whose computational cost typically grows rapidly with the number of calibrated parameters, the controller-based formulation introduces one additional constraint equation per parameter–target pair. As a result, expanding the calibrated parameter set does not fundamentally alter the stability of the optimisation landscape and incurs only marginal computational overhead.

In the present work, calibration was performed for individual steady-state snapshots. However, the proposed framework naturally extends to the calibration of longitudinal data, where successive time points are expected to yield closely related solutions. In this setting, the calibrated parameter set at a given time provides a natural initialisation for the subsequent calibration step, allowing the controllers to efficiently track gradual physiological changes and follow patient-specific trends over the course of an ICU stay.

## ACKNOWLEDGMENTS

The authors are grateful for the funding and support of the EPSRC-funded CHIMERA Maths in Healthcare Hub (EP/T017791/1), the Wellcome/EPSRC Centre for Interventional and Surgical Sciences (WEISS) (203145/A/16/Z), the Department of Mechanical Engineering and Department of Mathematics at UCL.

## DECLARATION OF GENERATIVE AI AND AI-ASSISTED TECHNOLOGIES IN THE WRITING PROCESS

During the preparation of this work the author(s) used GPT4 in order to Correct grammar and improve readability. After using this tool/service, the author(s) reviewed and edited the content as needed and take(s) full responsibility for the content of the publication.

**A APPENDIX**

## A.1 Nomenclature

**Table 6.**
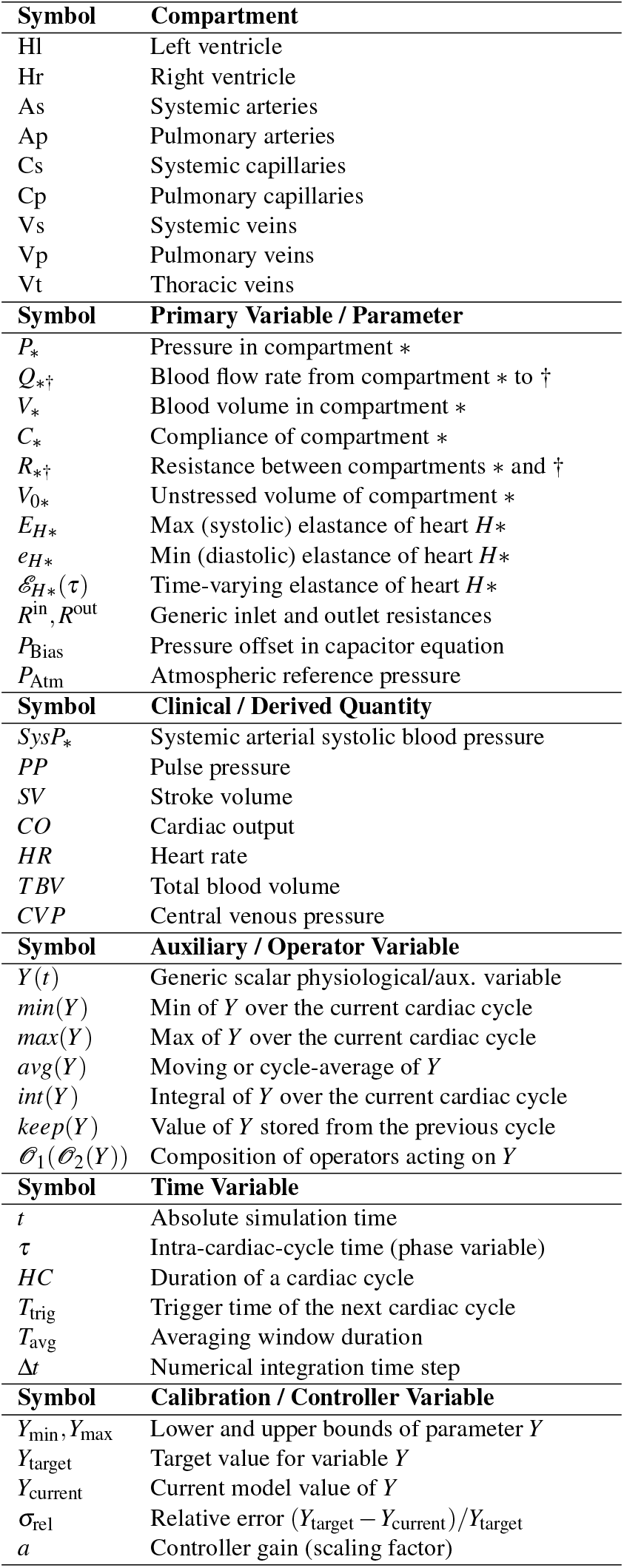
Nomenclature used in the cardiovascular model and calibration framework.

| Symbol | Compartment |
| --- | --- |
| Hl | Left ventricle |
| Hr | Right ventricle |
| As | Systemic arteries |
| Ap | Pulmonary arteries |
| Cs | Systemic capillaries |
| Cp | Pulmonary capillaries |
| Vs | Systemic veins |
| Vp | Pulmonary veins |
| Vt | Thoracic veins |
| Symbol | Primary Variable / Parameter |
| $P_*$ | Pressure in compartment * |
| $Q_{* \dagger}$ | Blood flow rate from compartment * to † |
| $V_*$ | Blood volume in compartment * |
| $C_*$ | Compliance of compartment * |
| $R_{* \dagger}$ | Resistance between compartments * and † |
| $V_{0*}$ | Unstressed volume of compartment * |
| $E_{H*}$ | Max (systolic) elastance of heart $H^*$ |
| $e_{H*}$ | Min (diastolic) elastance of heart $H^*$ |
| $\mathcal{E}_{H*}(\tau)$ | Time-varying elastance of heart $H^*$ |
| $R^{\text{in}}, R^{\text{out}}$ | Generic inlet and outlet resistances |
| $P_{\text{Bias}}$ | Pressure offset in capacitor equation |
| $P_{\text{Atm}}$ | Atmospheric reference pressure |
| Symbol | Clinical / Derived Quantity |
| $\text{Sys}P_*$ | Systemic arterial systolic blood pressure |
| $PP$ | Pulse pressure |
| $SV$ | Stroke volume |
| $CO$ | Cardiac output |
| $HR$ | Heart rate |
| $TBV$ | Total blood volume |
| $CVP$ | Central venous pressure |
| Symbol | Auxiliary / Operator Variable |
| $Y(t)$ | Generic scalar physiological/aux. variable |
| $\min(Y)$ | Min of $Y$ over the current cardiac cycle |
| $\max(Y)$ | Max of $Y$ over the current cardiac cycle |
| $\text{avg}(Y)$ | Moving or cycle-average of $Y$ |
| $\text{int}(Y)$ | Integral of $Y$ over the current cardiac cycle |
| $\text{keep}(Y)$ | Value of $Y$ stored from the previous cycle |
| $\mathcal{O}_1(\mathcal{O}_2(Y))$ | Composition of operators acting on $Y$ |
| Symbol | Time Variable |
| $t$ | Absolute simulation time |
| $\tau$ | Intra-cardiac-cycle time (phase variable) |
| $HC$ | Duration of a cardiac cycle |
| $T_{\text{trig}}$ | Trigger time of the next cardiac cycle |
| $T_{\text{avg}}$ | Averaging window duration |
| $\Delta t$ | Numerical integration time step |
| Symbol | Calibration / Controller Variable |
| $Y_{\min}, Y_{\max}$ | Lower and upper bounds of parameter $Y$ |
| $Y_{\text{target}}$ | Target value for variable $Y$ |
| $Y_{\text{current}}$ | Current model value of $Y$ |
| $\sigma_{\text{rel}}$ | Relative error $(Y_{\text{target}} - Y_{\text{current}})/Y_{\text{target}}$ |
| $a$ | Controller gain (scaling factor) |

## A.2 Model equations

### Capacitor Equations

The pressures for all capacitive elements listed in Table 3 follow the generic compliance relationship of Eq. 1. For each compartment *x*, the pressure is given by

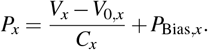

Using the symbols from the table, the corresponding expressions are:

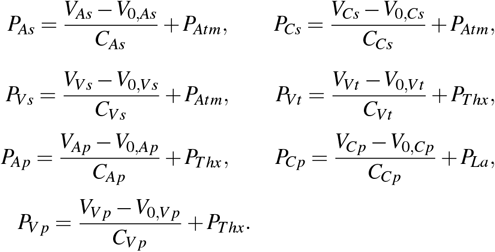

The ventricles use time-varying elastance (Eq. 7):

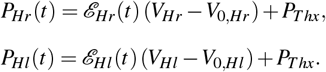

### Elastance Definitions

The time-varying elastance functions for the left and right heart chambers follow the generic formulation of Eq. 7, with chamber-specific parameters.

For the left heart (Hl), the elastance is defined as

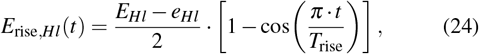

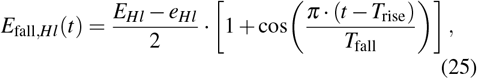

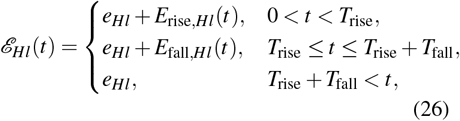

Similarly, for the right heart (Hr), the elastance is given by

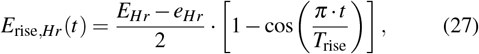

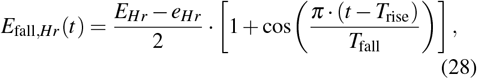

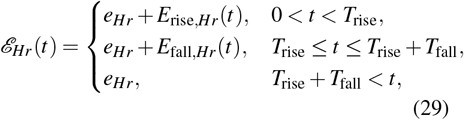

### Resistor and Diode Flow Equations

The resistive elements in Table 3 follow either the linear resistor law (Eq. 2)

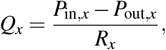

or the diode law (Eq. 4)

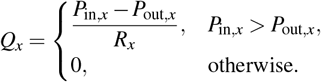

The symbolic flow expressions are:

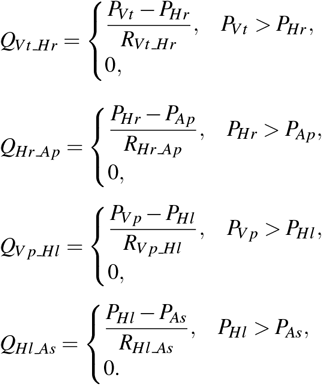

Linear resistors:

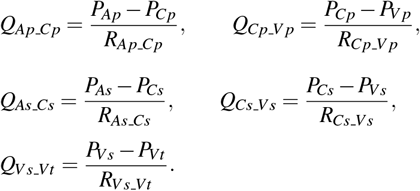

### Volume Balance Equations

All compartment volumes in Table 3 evolve according to the conservation law of Eq. 3:

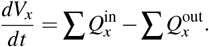

The symbolic volume dynamics are therefore:

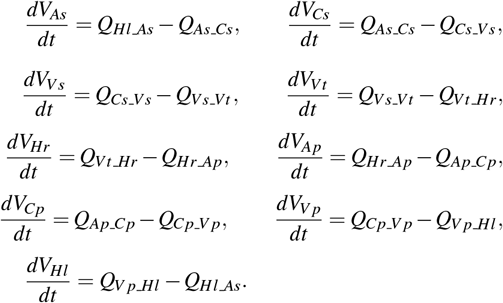

### Calibration variables based on cycle averages and integrals

The calibration targets in Table 2 are computed by instantiating the generic max/min, integral, keeper, and moving-average equations (12–15) with specific model variables. Below we write the explicit forms for the quantities used in the calibration procedure.

### Cycle-averaged compartment volumes

The cycle-averaged volumes are obtained using the moving-average equation 15, with *Y* (*T*) = *V*_*x*_(*T*) and *Avg*_*Y*_ (*T*) = *avg _V _x*(*T*):

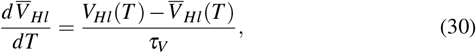

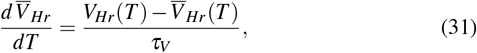

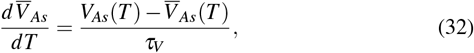

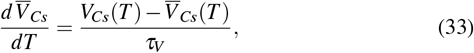

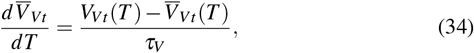

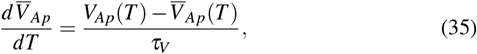

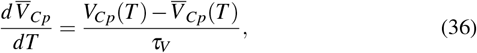

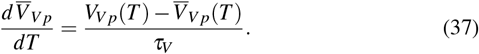

where *τ*_*V*_ is the averaging window (e.g. several seconds or beats). The systemic venous volume *V*_*Vs*_ is used directly as a state variable and not averaged.

### Stroke volume of the left ventricle

Stroke volume is computed from the flow *Q*_*Hl*_ *_*_*As*_ using the integral-reset equation 13, with *Y* (*T*) = *Q*_*Hl*_ *_*_*As*_(*T*):

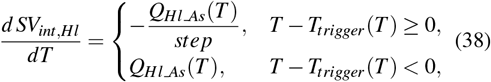

where *SV*_*int,Hl*_ accumulates the outflow over a single cardiac cycle. The cycle-wise stroke volume used for calibration, *keep _SV _Hl*, is obtained via the keeper equation 14 with *Y* (*T*) = *SV*_*int,Hl*_(*T*):

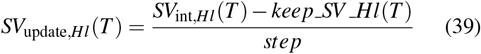

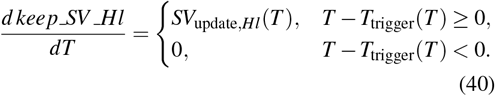

### Systolic, diastolic, and pulse pressures in the systemic arteries

For the systemic arterial pressure *P*_*As*_, we define within-cycle maxima and minima using the max/min equations (12–11) with *Y* (*T*) = *P*_*As*_(*T*):

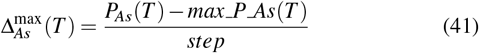

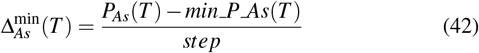

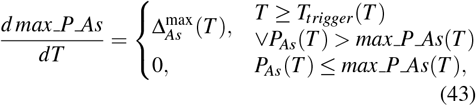

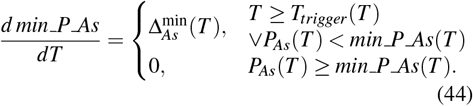

The cycle-wise systolic and diastolic pressures used in calibration are kept using Eq. 14:

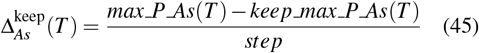

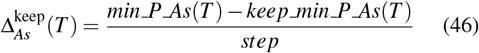

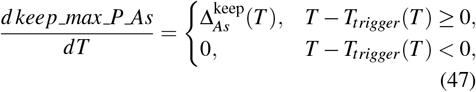

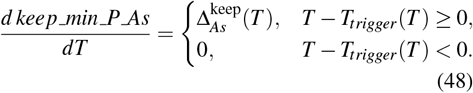

The systemic arterial pulse pressure variable appear

### Systolic, diastolic, and pulse pressures in the systemic arteries

For the systemic arterial pressure *P*_*As*_, within-cycle maxima and minima are computed using Eqs. (12–11) with *Y* (*T*) = *P*_*As*_(*T*).

#### Maxima and minima update terms

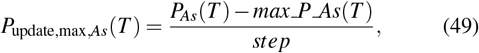

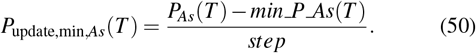

#### Within-cycle extrema dynamics

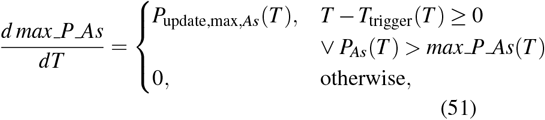

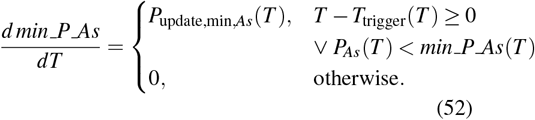

### Cycle-wise pressure retention

The cycle-wise systolic and diastolic pressures used for calibration are retained using Eq. 14.

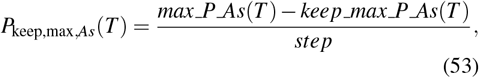

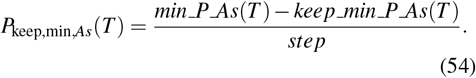

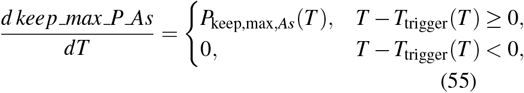

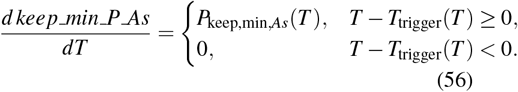

### Calibration controller equations

The calibration targets in Table 2 are enforced by embedding the model parameters as additional state variables and driving them with a cubic controller law. For each controller *j*, we define a dimensionless error *ε* _*j*_ between the model quantity *X*_*j*_(*T*) and its target value 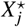,

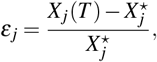

and update the associated parameter *θ*_*j*_(*T*) according to the cubic rule

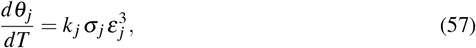

where *k* _*j*_ *>* 0 is a calibration gain and *σ*_*j*_ ∈ { +1, 1 } encodes the sign of the correlation in Table 4 (‘+’ for positive, ‘−’ for negative). A positive error *ε* _*j*_ *>* 0 therefore drives *θ*_*j*_ in the direction indicated by the correlation sign.

Below we list the explicit error definitions *X*_*j*_(*T*) and parameter dynamics *θ*_*j*_(*T*) for each controller used in Table 2. Superscripts ^⋆^ denote the corresponding target values (e.g. midpoints of the reported physiological ranges) for the selected phenotype (warm shock, cold shock, or normal).

**TBV controllers** Errors:

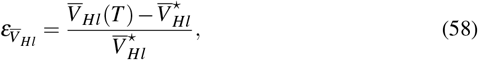

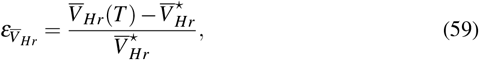

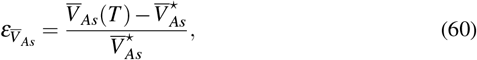

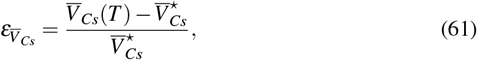

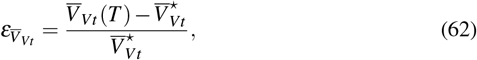

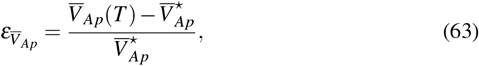

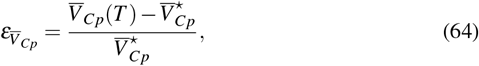

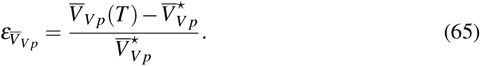

Parameter dynamics (correlation sign taken from Table 2):

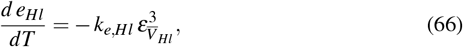

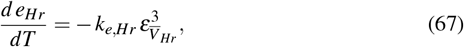

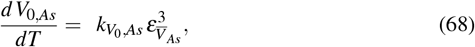

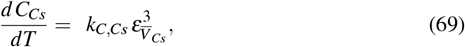

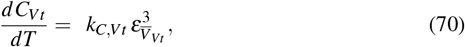

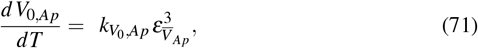

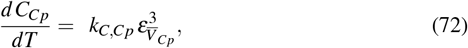

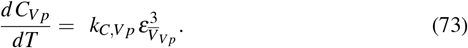

### Cardiac output and heart rate controller

Stroke volume of the left ventricle is obtained by cycle integration of *Q _Hl _As*(*T*) and stored as *keep _SV _Hl*. The corresponding error is

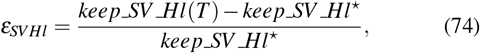

and the controller on the maximal left-ventricular elastance *E*_*Hl*_ is

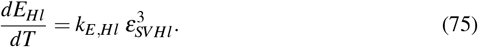

### Systemic arterial pressure and pulse pressure controllers

Let *keep _max _P _As* and *keep _min _P _As* denote the cycle-wise systolic and diastolic systemic arterial pressures. Their targets are *keep _max _P _As*^⋆^ and *amp _P _As*^⋆^ (pulse pressure), with

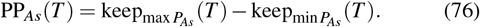

We define

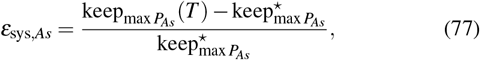

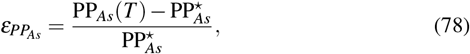

and drive the parameters

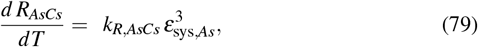

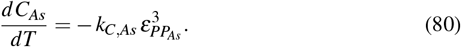

### Central venous pressure controller

For the systemic venous pressure *P*_*Vs*_, the cycle-averaged value 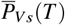 is compared with its target 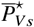:

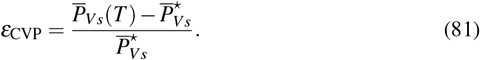

and the venous compliance controller is

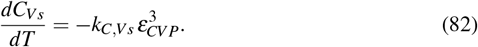

### Systemic capillary pressure controller

For the systemic capillary pressure *P*_*Cs*_, the cycle-averaged value 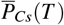 is used:

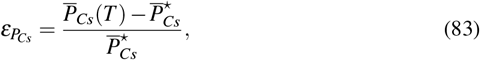

and the venous resistance is adjusted as

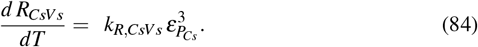

### Pulmonary arterial pressure controllers

For the pulmonary arterial pressure *P*_*Ap*_, we define cycle-wise systolic and diastolic values *keep _max _P*_*Ap*_(*T*) and *keep _min _P*_*Ap*_(*T*) with targets 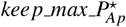 and 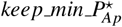:

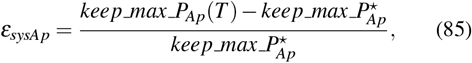

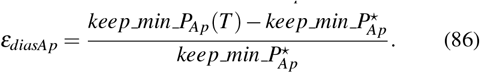

The corresponding controllers are

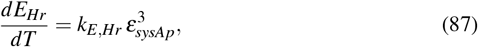

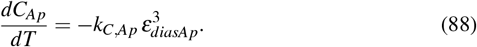

### Pulmonary capillary pressure controller

Finally, for the pulmonary capillary pressure *P*_*Cp*_, the cycle-averaged value 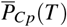 is compared with its target 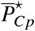:

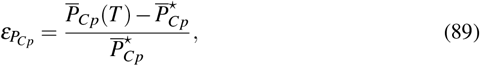

and the corresponding resistance controller is

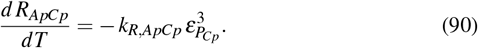

These equations instantiate the generic cubic controller law (16) for all calibration variables used in Table 2, linking each clinical target to a specific model parameter through the correlation structure indicated in the table.

## A.3 Calibration Example

**Figure 15.**
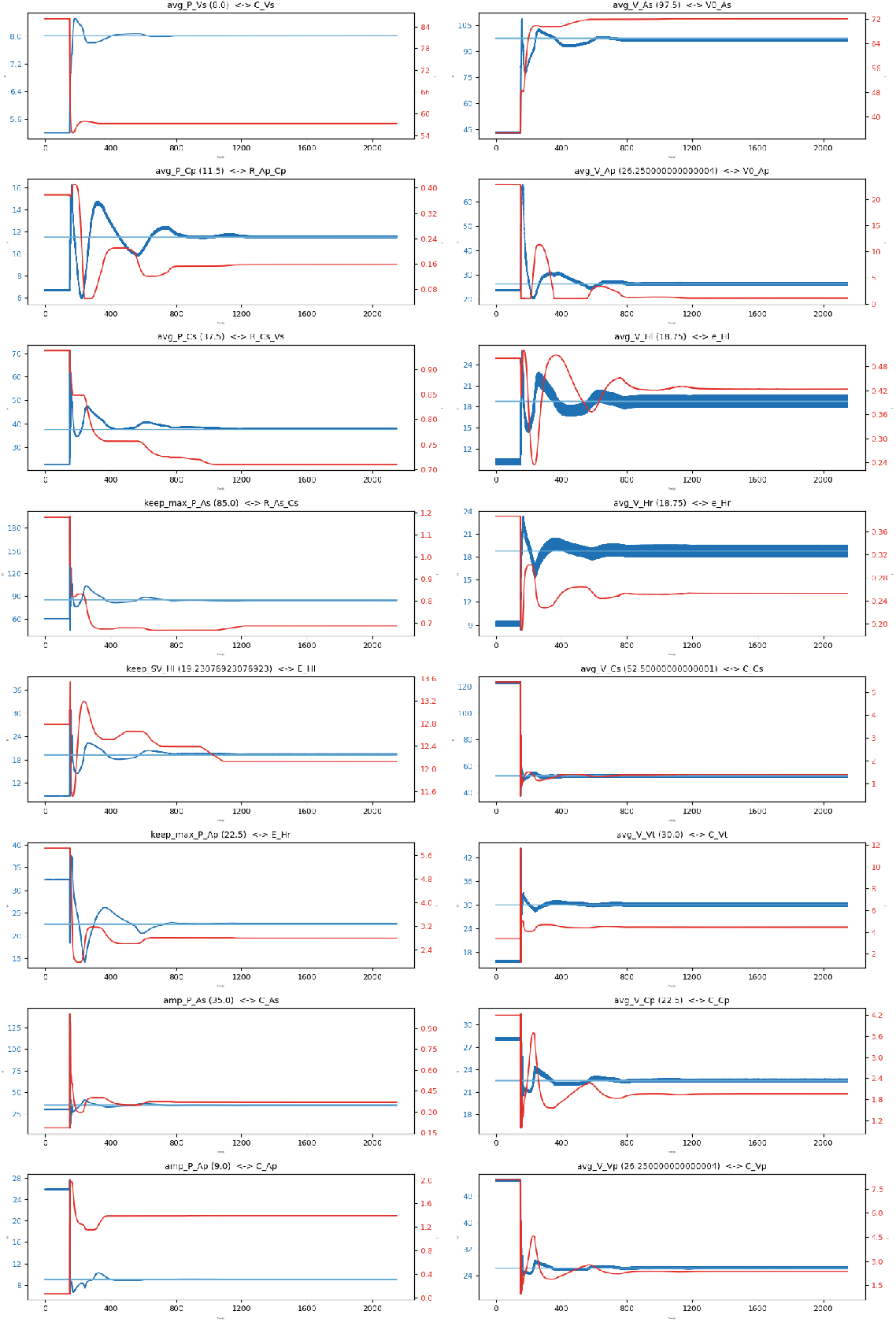
Representative calibration run showing the temporal evolution of selected model variables (blue), their targets (light blue), and the corresponding adaptive parameters (red). Parameter updates scale with the instantaneous deviation from the target and reverse sign upon crossing the target value. As equilibrium is approached, update magnitudes progressively diminish, resulting in stable bounded oscillations around the target.

## REFERENCES

[1] Krasimira Tsaneva-Atanasova and Vanessa Diaz-Zuccarini. Editorial: Mathematics for healthcare as part of computational medicine. Frontiers in Physiology, 9(JUL), 2018. ISSN 1664042X. doi: 10.3389/fphys.2018.00985.

[2] Mark Alber, Adrian Buganza Tepole, William R. Cannon, Suvranu De, Salvador Dura-Bernal, Krishna Garikipati, George Karniadakis, William W. Lytton, Paris Perdikaris, Linda Petzold, and Ellen Kuhl. Integrating machine learning and multiscale modeling—perspectives, challenges, and opportunities in the biological, biomedical, and behavioral sciences, 2019. ISSN 23986352.

[3] Nicolaas Westerhof, Frederik Bosman, Cornelis J. De Vries, and Abraham Noordergraaf. Analog studies of the human systemic arterial tree. Journal of Biomechanics, 2(2), 1969. ISSN 00219290. doi: 10.1016/0021-9290(69)90024-4.

[4] Catriona Stokes, Mirko Bonfanti, Zeyan Li, Jiang Xiong, Duanduan Chen, Stavroula Balabani, and Vanessa Díaz-Zuccarini. A novel MRI-based data fusion methodology for efficient, personalised, compliant simulations of aortic haemodynamics. Journal of Biomechanics, 129, 2021. ISSN 18732380. doi: 10.1016/j.jbiomech.2021.110793.

[5] Nico H.L. Kuijpers, Willem Dassen, Peter M. Van Dam, Eelco M. Van Dam, Evelien Hermeling, Joost Lumens, Theo Arts, and Tammo Delhaas. CircAdapt: A user-friendly learning environment for (patho) physiology of heart and circulation. In Computing in Cardiology, volume 39, 2012.

[6] Bram W. Smith, J. Geoffrey Chase, Roger I. Nokes, Geoffrey M. Shaw, and Graeme Wake. Minimal haemodynamic system model including ventricular interaction and valve dynamics. Medical Engineering and Physics, 26(2), 2004. ISSN 13504533. doi: 10.1016/j.medengphy.2003.10.001.

[7] Bram W. Smith, Steen Andreassen, Geoffrey M. Shaw, Per L. Jensen, Stephen E. Rees, and J. Geoffrey Chase. Simulation of cardiovascular system diseases by including the autonomic nervous system into a minimal model. Computer Methods and Programs in Biomedicine, 86(2), 2007. ISSN 01692607. doi: 10.1016/j.cmpb.2007.02.001.

[8] Thomas Heldt, Eun B. Shim, Roger D. Kamm, Roger G. Mark, and Massachusetts. Computational modeling of cardiovascular response to orthostatic stress. Journal of Applied Physiology, 92(3), 2002. ISSN 87507587. doi: 10.1152/japplphysiol.00241.2001.

[9] Sonia Charleston-Villalobos, Sina Reulecke, Andreas Voss, Mahmood R. Azimi-Sadjadi, Ramón González-Camarena, Mercedes J. Gaitán-González, Jesús A. González-Hermosillo, Guadalupe Hernández-Pacheco, Steffen Schulz, and Tomás Aljama-Corrales. Time-frequency analysis of cardiovascular and cardiorespiratory interactions during orthostatic stress by extended partial directed coherence. Entropy, 21(5), 2019. ISSN 10994300. doi: 10.3390/e21050468.

[10] MT Cabeleira, DV Anand, S Ray, C Black, C C Ovenden, and V Díaz-Zuccarini. Comparing physiological impacts of positive pressure ventilation versus self-breathing via a versatile cardiopulmonary model incorporating a novel alveoli opening mechanism. Computers in Biology and Medicine, 180:108960, 2024. ISSN 0010-4825. doi: 10.1016/j.compbiomed.2024.108960. URL https://www.sciencedirect.com/science/article/pii/S001048252401045X.

[11] K. Lu, J. W. Clark, F. H. Ghorbel, D. L. Ware, and A. Bidani. A human cardiopulmonary system model applied to the analysis of the Valsalva maneuver. American Journal of Physiology - Heart and Circulatory Physiology, 281(6 50-6), 2001. ISSN 03636135. doi: 10.1152/ajpheart.2001.281.6.h2661.

[12] Elisa Magosso and Mauro Ursino. A mathematical model of CO2 effect on cardiovascular regulation. American Journal of Physiology - Heart and Circulatory Physiology, 281(5 50-5), 2001. ISSN 03636135. doi: 10.1152/ajpheart.2001.281.5.h2036.

[13] Chuong Ngo, Stephan Dahlmanns, Thomas Vollmer, Berno Misgeld, and Steffen Leonhardt. An object-oriented computational model to study cardiopulmonary hemodynamic interactions in humans. Computer Methods and Programs in Biomedicine, 159, 2018. ISSN 18727565. doi: 10.1016/j.cmpb.2018.03.008.

[14] A. Albanese, N. W. Chbat, and M. Ursino. Transient respiratory response to hypercapnia: Analysis via a cardiopulmonary simulation model. In Proceedings of the Annual International Conference of the IEEE Engineering in Medicine and Biology Society, EMBS, 2011. doi: 10.1109/IEMBS.2011.6090668.

[15] Antonio Albanese, Limei Cheng, Mauro Ursino, and Nicolas W. Chbat. An integrated mathematical model of the human cardiopulmonary system: Model development. American Journal of Physiology - Heart and Circulatory Physiology, 310 (7), 2016. ISSN 15221539. doi: 10.1152/ajpheart.00230.2014.

[16] Luciano G. Fernandes, Paulo R. Trenhago, Raúl A. Feijóo, and Pablo J. Blanco. Integrated cardiorespiratory system model with short timescale control mechanisms. International Journal for Numerical Methods in Biomedical Engineering, 37 (11), 2021. ISSN 20407947. doi: 10.1002/cnm.3332.

[17] Gustavo Guerrero, Virginie Le Rolle, and Alfredo Hernández. Parametric Analysis of an Integrated Model of Cardio-respiratory Interactions in Adults in the Context of Obstructive Sleep Apnea. Annals of Biomedical Engineering, 49(12), 2021. ISSN 15739686. doi: 10.1007/s10439-021-02828-6.

[18] X Liu, C Mo, J Li, H Yu, S Hu, d. Zhu, P Zhang, and Y Li. Development of a Lumped Parameter Model of Human Whole Body Circulatory Loop. IEEE Access, 12:188505–188518, 2024. ISSN 2169-3536. doi: 10.1109/ACCESS.2024.3491112.

[19] Neta Ravid Tannenbaum, Omer Gottesman, Azadeh Assadi, Mjaye Mazwi, Uri Shalit, and Danny Eytan. iCVS-Inferring Cardio-Vascular hidden States from physiological signals available at the bedside. PLoS Computational Biology, 19(9 September), 2023. ISSN 15537358. doi: 10.1371/journal.pcbi.1010835.

[20] Yekanth Ram Chalumuri, Ghazal Arabidarrehdor, Ali Tivay, Catherine M. Sampson, Muzna Khan, Michael Kinsky, George C. Kramer, Jin-Oh Hahn, Christopher G. Scully, and Ramin Bighamian. A Lumped-Parameter Model of the Cardiovascular System Response for Evaluating Automated Fluid Resuscitation Systems. IEEE Access, 12:62511–62525, 2024. doi: 10.1109/ACCESS.2024.3395008.

[21] R. T. Djoumessi, I. P. Dongmo Vougmo, J. S. Tadjonang Tegne, and F. B. Pelap. Proposed cardio-pulmonary model to investigate the effects of COVID-19 on the cardiovascular system. Heliyon, 9(1), 2023. ISSN 24058440. doi: 10.1016/j.heliyon.2023.e12908.

[22] James Cushway, Liam Murphy, J. Geoffrey Chase, Geoffrey M. Shaw, and Thomas Desaive. Physiological trend analysis of a novel cardio-pulmonary model during a preload reduction manoeuvre. Computer Methods and Programs in Biomedicine, 220, 2022. ISSN 18727565. doi: 10.1016/j.cmpb.2022.106819.

[23] Harry Saxton, Xu Xu, Torsten Schenkel, and Ian Halliday. Assessing input parameter hyperspace and parameter identifiability in a cardiovascular system model via sensitivity analysis. Journal of Computational Science, 79:102287, 7 2024. ISSN 1877-7503. doi: 10.1016/J.JOCS.2024.102287.

[24] Finbar Argus, Debbie Zhao, Thiranja P. Babarenda Gamage, Martyn P. Nash, and Gonzalo D. Maso Talou. Automated model calibration with parallel MCMC: Applications for a cardiovascular system model. Frontiers in Physiology, 13, 2022. ISSN 1664042X. doi: 10.3389/fphys.2022.1018134.

[25] Bálint Szabó, Á kos Antal, Á kos Szlávecz, Béla Paláncz, Katalin Kovács, Liam Murphy, James Cushway, Nicolas Davey, Cong Zhou, J. Geoffrey Chase, and Balázs Benyó. Cardiovascular Model Identification Using Neural ODE. IFAC-PapersOnLine, 58(24):374–379, 1 2024. ISSN 2405-8963. doi: 10.1016/J.IFACOL.2024.11.066.

[26] Wolfgang J. Kern, Simon Orlob, Gabriel Putzer, Judith Martini, and Martin Holler. A Parameter Identification Approach towards Analyzing Hemodynamics Based on Capnography. In Computing in Cardiology, 2023. doi: 10.22489/CinC.2023.086.

[27] Mitchel J. Colebank, Pim A. Oomen, Colleen M. Witzenburg, Anna Grosberg, Daniel A. Beard, Dirk Husmeier, Mette S. Olufsen, and Naomi C. Chesler. Guidelines for mechanistic modeling and analysis in cardiovascular research, 2024. ISSN 15221539.

[28] Luregn J. Schlapbach, R. Scott Watson, Lauren R. Sorce, Andrew C. Argent, Kusum Menon, Mark W. Hall, Samuel Akech, David J. Albers, Elizabeth R. Alpern, Fran Balamuth, Melania Bembea, Paolo Biban, Enitan D. Carrol, Kathleen Chiotos, Mohammod Jobayer Chisti, Peter E. Dewitt, Idris Evans, Cláudio Flauzino De Oliveira, Christopher M. Horvat, David Inwald, Paul Ishimine, Juan Camilo Jaramillo-Bustamante, Michael Levin, Rakesh Lodha, Blake Martin, Simon Nadel, Satoshi Nakagawa, Mark J. Peters, Adrienne G. Randolph, Suchitra Ranjit, Margaret N. Rebull, Seth Russell, Halden F. Scott, Daniela Carla De Souza, Pierre Tissieres, Scott L. Weiss, Matthew O. Wiens, James L. Wynn, Niranjan Kissoon, Jerry J. Zimmerman, L. Nelson Sanchez-Pinto, and Tellen D. Bennett. International Consensus Criteria for Pediatric Sepsis and Septic Shock. JAMA, 331(8), 2024. ISSN 15383598. doi: 10.1001/jama.2024.0179.

[29] Mervyn Singer, Clifford S. Deutschman, Christopherwarren Seymour, Manu Shankar-Hari, Djillali Annane, Michael Bauer, Rinaldo Bellomo, Gordon R. Bernard, Jean Daniel Chiche, Craig M. Coopersmith, Richard S. Hotchkiss, Mitchell M. Levy, John C. Marshall, Greg S. Martin, Steven M. Opal, Gordon D. Rubenfeld, Tomvan Der Poll, Jean Louis Vincent, and Derek C. Angus. The third international consensus definitions for sepsis and septic shock (sepsis-3), 2 2016. ISSN 15383598.

[30] Manu Shankar-Hari, Gary S. Phillips, Mitchell L. Levy, Christopher W. Seymour, Vincent X. Liu, Clifford S. Deutschman, Derek C. Angus, Gordon D. Rubenfeld, and Mervyn Singer. Developing a newdefinition and assessing newclinical criteria for Septic shock: For the third international consensus definitions for sepsis and septic shock (sepsis-3). JAMA - Journal of the American Medical Association, 315(8), 2016. ISSN 15383598. doi: 10.1001/jama.2016.0289.

[31] Luca Marchetto, Marco Daverio, Rosanna Comoretto, Davide Padrin, Serena Scaravetti, Giulia Bordin, Stefania Ferrario, Maria Cristina Mondardini, Enzo Picconi, Immacolata Rulli, Francesco Sacco, Pasquale Vitale, Gloria Brigiari, Luregn J Schlapbach, Kusum Menon, Dario Gregori, Angela Amigoni, and for the Italian Network of PICU Study Group (TIPNet). Predictive and Prognostic Performance of the Phoenix Sepsis Criteria and Phoenix Sepsis Score in PICU Patients With Suspected Infection: A Multicenter Prospective Study. Critical Care Medicine, 2026. ISSN 1530-0293.

[32] Scott L. Weiss, Mark J. Peters, Waleed Alhazzani, Michael S.D. Agus, Heidi R. Flori, David P. Inwald, Simon Nadel, Luregn J. Schlapbach, Robert C. Tasker, Andrew C. Argent, Joe Brierley, Joseph Carcillo, Enitan D. Carrol, Christopher L. Carroll, Ira M. Cheifetz, Karen Choong, Jeffry J. Cies, Andrea T. Cruz, Daniele De Luca, Akash Deep, Saul N. Faust, Claudio Flauzino De Oliveira, Mark W. Hall, Paul Ishimine, Etienne Javouhey, Koen F.M. Joosten, Poonam Joshi, Oliver Karam, Martin C.J. Kneyber, Joris Lemson, Graeme MacLaren, Nilesh M. Mehta, Morten Hylander Møller, Christopher J.L. Newth, Trung C. Nguyen, Akira Nishisaki, Mark E. Nunnally, Margaret M. Parker, Raina M. Paul, Adrienne G. Randolph, Suchitra Ranjit, Lewis H. Romer, Halden F. Scott, Lyvonne N. Tume, Judy T. Verger, Eric A. Williams, Joshua Wolf, Hector R. Wong, Jerry J. Zimmerman, Niranjan Kissoon, and Pierre Tissieres. Surviving sepsis campaign international guidelines for the management of septic shock and sepsis-associated organ dysfunction in children. Intensive Care Medicine, 46, 2020. ISSN 14321238. doi: 10.1007/s00134-019-05878-6.

[33] Mariana Miranda and Simon Nadel. Pediatric Sepsis: a Summary of Current Definitions and Management Recommendations. Current Pediatrics Reports, 11(2):29–39, 2023. ISSN 2167-4841. doi: 10.1007/s40124-023-00286-3. URL 10.1007/s40124-023-00286-3.

[34] Akash Deep, Chulananda D.A. Goonasekera, Yanzhong Wang, and Joe Brierley. Evolution of haemodynamics and outcome of fluid-refractory septic shock in children. Intensive Care Medicine, 39(9), 2013. ISSN 03424642. doi: 10.1007/s00134-013-3003-z.

[35] R. K. Aneja and J. A. Carcillo. Differences between adult and pediatric septic shock, 2011. ISSN 03759393.

[36] Kathryn Maitland, Sarah Kiguli, Robert O. Opoka, Charles Engoru, Peter Olupot-Olupot, Samuel O. Akech, Richard Nyeko, George Mtove, Hugh Reyburn, Trudie Lang, Bernadette Brent, Jennifer A. Evans, James K. Tibenderana, Jane Crawley, Elizabeth C. Russell, Michael Levin, Abdel G. Babiker, and Diana M. Gibb. Mortality after Fluid Bolus in African Children with Severe Infection. New England Journal of Medicine, 364(26), 2011. ISSN 0028-4793. doi: 10.1056/nejmoa1101549.

[37] Susanna Esposito, Benedetta Mucci, Eleonora Alfieri, Angela Tinella, and Nicola Principi. Advances and Challenges in Pediatric Sepsis Diagnosis: Integrating Early Warning Scores and Biomarkers for Improved Prognosis, 2025. ISSN 2218273X.

[38] Ranaa Akkawi El Edelbi, Synnö ve Lindemalm, Per Nydert, and Staffan Eksborg. Estimation of body surface area in neonates, infants, and children using body weight alone. International Journal of Pediatrics and Adolescent Medicine, 8 (4), 2021. ISSN 23526467. doi: 10.1016/j.ijpam.2020.09.003.

[39] Ann Raes, Sarah Van Aken, Margarita Craen, Raymond Donckerwolcke, and Johan Vande Walle. A reference frame for blood volume in children and adolescents. BMC Pediatrics, 6, 2006. ISSN 14712431. doi: 10.1186/1471-2431-6-3.

[40] SeyedAli Khonsary. Guyton and Hall: Textbook of Medical Physiology. Surgical Neurology International, 8(1), 2017. ISSN 2152-7806. doi: 10.4103/sni.sni_327_17.

[41] Joe Brierley and Mark J. Peters. Distinct hemodynamic patterns of septic shock at presentation to pediatric intensive care. Pediatrics, 122(4), 2008. ISSN 00314005. doi: 10.1542/peds.2007-1979.

[42] H.D. O’Reilly and K. Menon. Sepsis in paediatrics, 2021. ISSN 20585357.

[43] Suchitra Ranjit and Rajeswari Natraj. Hemodynamic Management Strategies in Pediatric Septic Shock: Ten Concepts for the Bedside Practitioner, 2024. ISSN 09747559.

[44] Seung Jun Choi, Eun Ju Ha, Won Kyoung Jhang, and Seong Jong Park. Elevated central venous pressure is associated with increased mortality in pediatric septic shock patients. BMC Pediatrics, 18(1), 2018. ISSN 14712431. doi: 10.1186/s12887-018-1059-1.

[45] Blerina Asllanaj, Elizabeth Benge, Jieun Bae, and Yi McWhorter. Fluid management in septic patients with pulmonary hypertension, review of the literature, 2023. ISSN 2297055X.

[46] Bassam M. Gebara, Brahm Goldstein, Brett Giroir, and Adrienne Randolph. Values for systolic blood pressure, 2005. ISSN 15297535.

[47] Shane M. Tibby, Mark Hatherill, Michael J. Marsh, and Ian A. Murdoch. Clinicians’ abilities to estimate cardiac index in ventilated children and infants. Archives of Disease in Childhood, 77(6), 1997. ISSN 14682044. doi: 10.1136/adc.77.6.516.

[48] Sujata Deshpande, Pradeep Suryawanshi, Shrikant Holkar, Yogen Singh, Rameshwor Yengkhom, Jan Klimek, and Samir Gupta. Pulmonary hypertension in late onset neonatal sepsis using functional echocardiography: a prospective study. Journal of Ultrasound, 25(2), 2022. ISSN 18767931. doi: 10.1007/s40477-021-00590-y.

[49] Steven H. Abman, Georg Hansmann, Stephen L. Archer, D. Dunbar Ivy, Ian Adatia, Wendy K. Chung, Brian D. Hanna, Erika B. Rosenzweig, J. Usha Raj, David Cornfield, Kurt R. Stenmark, Robin Steinhorn, Bernard Thébaud, Jeffrey R. Fineman, Titus Kuehne, Jeffrey A. Feinstein, Mark K. Friedberg, Michael Earing, Robyn J. Barst, Roberta L. Keller, John P. Kinsella, Mary Mullen, Robin Deterding, Thomas Kulik, George Mallory, Tilman Humpl, and David L. Wessel. Pediatric pulmonary hypertension, 2015. ISSN 15244539.

